# A domesticated totivirus-like tandem array undergoes interspecific transfer and asymmetric evolution

**DOI:** 10.64898/2026.05.24.726934

**Authors:** Derek J. Taylor, Dominic A. Tringali

## Abstract

RNA paleoviruses are expected to evolve more slowly than their exogenous viral progenitors. We show that a four-gene tandem array (STORM, Scheffersomyces Totivirus-like Responsive Module; genes TLC1-TLC4) in wood-associated yeasts violates this expectation, evolving faster at the protein level than its exogenous totiviral relatives while persisting for over 15 million years. STORM has accumulated greater amino-acid divergence than its exogenous totiviral relatives over a much shorter host phylogenetic window (∼54 MY of *Scheffersomyces* history versus ∼225 MY for exogenous totivirus diversification), under significant relaxation of selective constraint (RELAX K < 1). Tandem duplication resulted in asymmetric evolution within the array. For example, TLC4 alone has retained the predicted decapping loop motif (lost from TLC1, TLC2, and TLC3) and a totivirus-like capsid fold. Other copies remain more constrained in structure and sequence, indicating functional partitioning. All four genes are transcriptionally active, embedded in host antiviral and RNA-decay regulatory neighborhoods, with condition-dependent expression. Hundreds of reference gene trees for *Scheffersomyces* are concordant with the species tree, with only two unrelated singleton exceptions; the STORM array is the only locus where all paralogs share a well-supported, locus-coherent discordance. Distance-based tests are inconsistent with incomplete lineage sorting, and shared discordance with an adjacent ATP10 pseudogene and a transposase (Tc1/mariner superfamily) implicates transposon-mediated co-mobilization. We infer at least two interspecific transfers of STORM. Our results reveal how hosts can domesticate a mobile virus-like module whose paralogs escape strong purifying selection and explore sequence space while the core fold is conserved.

## Introduction

Over decades to centuries, the rapid evolution of many RNA viruses is readily observed (Duffy, et al. 2008; Duchêne, et al. 2014; Mifsud, et al. 2025). However, the use of outbreak-calibrated molecular clocks often leads to vastly younger divergence time estimates than the minimum ages inferred from the ancient paleoviral record and dated host genomes (Simmonds, et al. 2019). Paleoviruses arise when viral genetic material becomes integrated and retained in host genomes (Patel, et al. 2011). One explanation for this discrepancy is that rapid short-term viral evolution produces extensive substitution saturation over longer timescales, especially in compact genomes with constrained proteins. As a result, apparent long-term rates decline and can approach host-like evolutionary rates. The Prisoner-of-War (PoW) model formalizes this time-dependent rate decay by modeling the transition from rapidly evolving viral regimes to more constrained long-term evolutionary dynamics (Ghafari, et al. 2021; Ghafari, et al. 2024). Here we report a domesticated tandem viral array in yeast that appears to defy the expected post-integration slowdown.

Although ancient paleoviruses helped to motivate the development of time-dependent rate models, they themselves remain under-explored as evolutionary systems. Paleoviruses potentially represent natural rate-switch experiments, with a pre-integration viral stage followed by a post-integration host stage. They can confer novel functions, provide fossil-like anchors for deep-time calibrations, and illuminate historical host ranges (Holmes 2011; Patel, et al. 2011; Taylor, et al. 2014; Ghosh, et al. 2024; Barreat, et al. 2025; Hu, et al. 2025). Duplication of non-retroviral paleoviral loci has been observed, including tandem expansions in some lineages (Ballinger, et al. 2014). Although little is known about tandem paleoviruses, tandem duplication in general can boost expression and drive asymmetric divergence among paralogs, as redundant copies relax purifying selection on individual members (Loehlin and Carroll 2016; Holland, et al. 2017; Fan, et al. 2023). The rate acceleration can sometimes be so severe as to obscure phylogenetic reconstruction, with proposed mechanisms including rewiring of regulatory regions followed by neofunctionalization, resulting in conserved and exploratory copies. Tandem paleoviral arrays are particularly tractable for rate inference: their members are expected to share a single integration ancestor, whereas dispersed singleton paleoviruses may reflect multiple independent integrations and cannot be assumed to share a common origin.

Totiviruses are dsRNA viruses that primarily infect fungi (e.g., yeasts and smut fungi) and are transmitted mainly through cell division, although occasional host jumps are inferred from phylogenetic evidence including a plant-associated clade (Hughes, et al. 2024).

Nonretroviral RNA paleoviruses related to totiviruses are duplicated in yeast (Taylor and Bruenn 2009). Yeast-associated L-A and L-BC viruses snatch the caps of host mRNAs (via a decapping structure; Fujimura and Esteban 2011), exposing transcripts to degradation by the 5′→3′ exonuclease XRN1 and the associated SKI (superkiller) antiviral complex (Masison, et al. 1995). In *Saccharomyces*, totiviruses can impose fitness costs (e.g. via proteotoxic stress) but also may confer benefits, such as improved responses to environmental stress (Hsiao, et al. 2025). A tight co-evolutionary relationship is evident between totiviruses and host RNA decay machinery, with yeast expressing both strong antiviral factors (e.g. SKI3) and virus-maintaining factors (Masison, et al. 1995; Rowley, et al. 2016; Chau, et al. 2023; Hsiao, et al. 2025).

The genome of *Scheffersomyces stipitis* (Serinales; strain CBS 6054) harbors a unique capsid-like gene array related to L-A totivirus (Taylor and Bruenn 2009). The genome of *S. stipitis* includes a complete polyprotein with RNA-dependent RNA polymerase (RdRp) and capsid-like regions plus three additional capsid-like paralogs, each preserving a continuous open reading frame. A protein product has been demonstrated for one capsid-like paralog in *S. stipitis* (Taylor, et al. 2013), and the heterogeneity in gene architecture (from a full polyprotein to capsid-only copies) suggests asymmetric constraint among duplicates. Indeed, earlier phylogenetic analyses with only one STORM-containing genome failed to resolve the phylogenetic position of capsid-like paleoviruses of *S. stipitis* (strain CBS 6054) (Taylor and Bruenn 2009; Brejova, et al. 2024).

Here we investigate the tempo and mode of evolution of totivirus-like genes across publicly available genomes of yeast from Serinales, focusing on wood- and insect-associated species of *Scheffersomyces*, which share ecological habitats conducive to interspecific contact (Shelomi 2025). We present an unusual case of a domesticated RNA viral array whose long-term protein evolution exceeds that of related exogenous viruses. We test three predictions for an asymmetric paleoviral module. First, if tandem duplication promotes asymmetric divergence, then one or more tandem paralogs should exhibit elevated amino-acid substitution rates and signatures of relaxed selection relative to more constrained copies and to related extant totiviruses. Second, if the tandem array has been integrated into host biology, then array organization, inferred protein structure, and transcriptional responsiveness should show conserved patterns across taxa. Third, if the paleoviral tandem array behaves as a long-lived mobile module capable of interspecific transfer, then its copies should retain evidence of shared ancestry as an array, while showing phylogenetic or genomic-context discordance inconsistent with simple vertical inheritance.

## Results

### Totivirus-like tandem paleoviral arrays in *Scheffersomyces*

Genome screening of Serinales yeasts identified totivirus-like capsid and RdRp domains in multiple *Scheffersomyces* genomes, but only a subset of species carried a distinctive four-gene tandem paleoviral array (Fig. 1): one polyprotein ORF (N-terminal capsid + C-terminal RdRp) plus three capsid-only ORFs. We refer to this array as STORM (Scheffersomyces Totivirus-like Responsive Module). The tandem totivirus-like capsid (TLC) paralogs are designated TLC1-TLC4 according to their order in the *S. stipitis* array, with TLC2 representing the capsid region of the capsid-RdRp polyprotein. Comparative gene prediction and ORF calling across species indicated largely conserved intronless ORF boundaries among array members. However, several species (*S. segobiensis* NRRL Y-11571, *S. quercinus*, *S. virginianus*, and *S. illinoinensis*) lacking the full four-gene array retained one or two fragmented, pseudogenized TLC-like remnants, consistent with independent decay following loss of an ancestral array.

**Figure 1.**
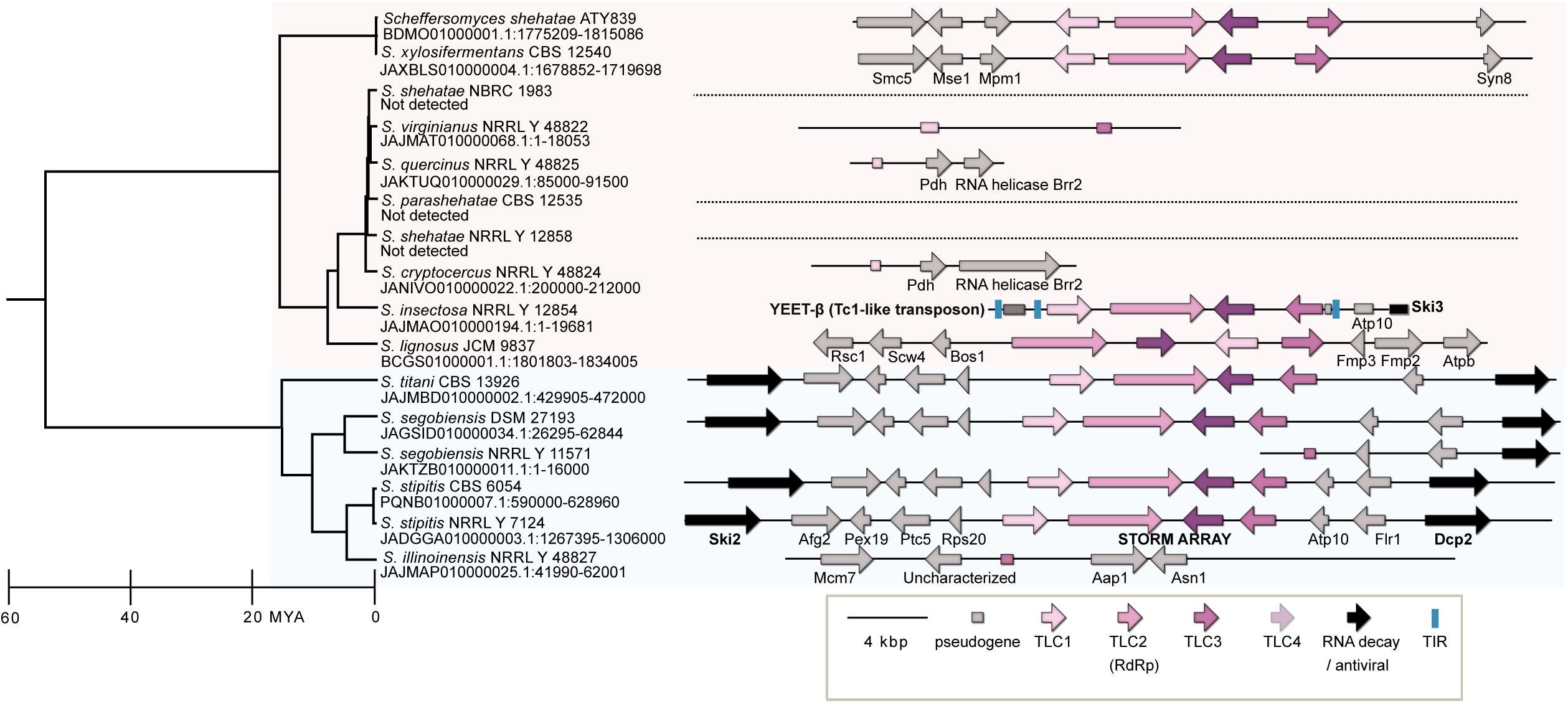
Time-calibrated phylogeny of *Scheffersomyces* and genomic contexts of the STORM paleoviral array. Left, chronogram for species of *Scheffersomyces* based on concatenated nuclear genes, with the time scale in millions of years before present (Mya). Right, gene neighborhoods surrounding the STORM array (colored blocks) mapped onto the tips of the tree; shaded rectangles represent the major clades of *Scheffersomyces* examined here. Arrows show ORFs and their transcriptional orientation; STORM components (TLC1-TLC4 and the polyprotein-like capsid-RdRp) are colored according to the legend (bottom). Selected neighboring host genes implicated in oxidative phosphorylation (OXPHOS) or antiviral function (e.g. SKI3, ATP10, DCP2) are labeled for each context. The architecture of a STORM-associated DNA transposon, YEET (Tc1-family), is shown for *S. insectosa*. Dotted lines mark genomes where the syntenic region is present, but the array has been lost or reduced to undetected fragments.

### Long-term conservation and rapid loss of array architecture

Microsynteny analysis using flanking regions (10-15 kb on either side) showed that the four-gene architecture is conserved among *Scheffersomyces* species in which it occurs (Fig. 1). Within the *titani*/*stipitis* clade, arrays occupy syntenic positions relative to neighboring host genes and maintain a conserved internal order (TLC1 → TLC2 [polyprotein] → TLC3 → TLC4). In the *lignosus* clade, arrays retain a STORM-like architecture but occur in at least three distinct chromosomal contexts, suggesting translocation and/or horizontal transfer events (Figs. 1-2).

**Figure 2.**
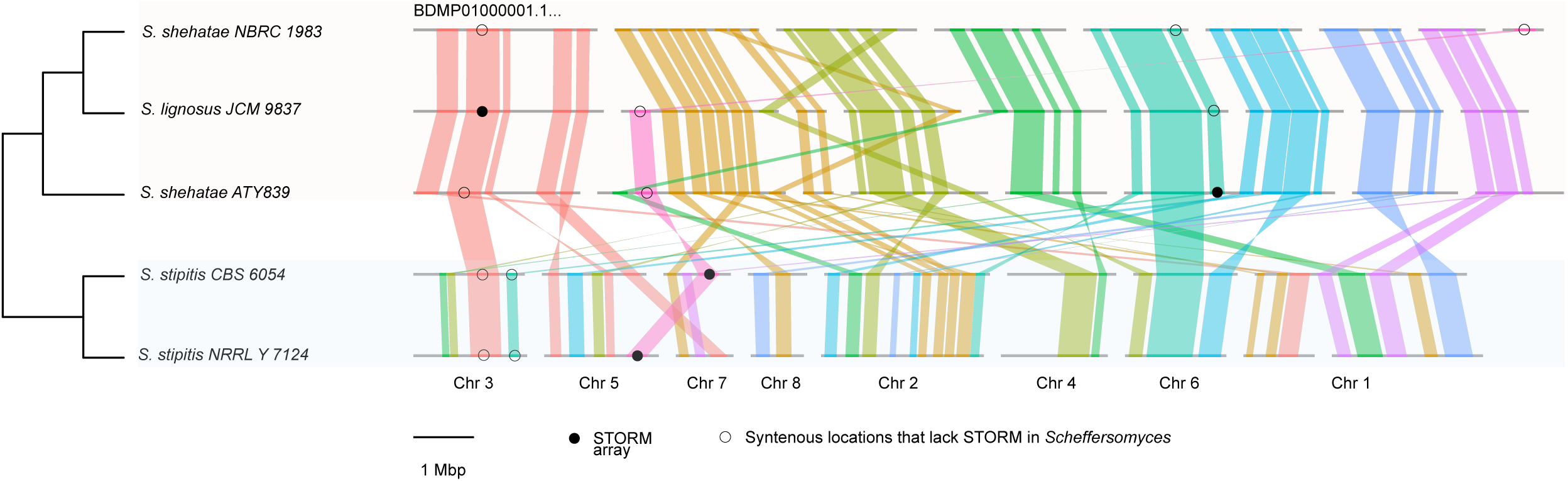
Macrosynteny and chromosomal locations of the STORM paleoviral array in Scheffersomyces. Macrosynteny plot (ntSynt) showing whole-genome alignments for five genomes of *Scheffersomyces* ordered according to the species tree on the left (shaded colors represent two major clades of wood-digesting species). Each horizontal gray bar represents a chromosome or major scaffold, with colored blocks and ribbons connecting syntenic segments among genomes. Filled circles mark the positions of intact STORM arrays, whereas open circles indicate syntenic locations in other genomes. In *S. stipitis* CBS 6054 and *S. stipitis* NRRL Y-7124 the arrays map to different chromosomes (Chr 7 and Chr 5, respectively). The bottom track shows the chromosome numbering for *S. stipitis* CBS 6054 (based on size), and the scale bar indicates 1 Mbp.

*S. lignosus* is the only species in our sampling in which TLC1 occurs internally within the array. Time-calibrated *Scheffersomyces* species trees based on concatenated nuclear genes place the common ancestor of the syntenic *titani*-clade array at more than 15 million years old (Fig. 1), implying long-term persistence of the four-gene module. The *S. lignosus* array is likely substantially older because *S. lignosus* diverged from *S. titani* approximately 43.2-65.1 MYA, and each *S. lignosus* TLC paralog exceeds the total divergence observed across the syntenic *titani*-clade arrays. Genomes lacking STORM arrays generally retain syntenic flanks with either a gap at the array position or decaying capsid-like fragments.

At broader chromosomal scale, ntSynt macrosynteny analyses revealed extensive rearrangements among deeper *Scheffersomyces* clades, yet STORM remains embedded within locally syntenic genomic neighborhoods (Fig. 2). Some insertions lie in subtelomeric regions, but we observed at least five distinct genomic contexts (including variation among *S. stipitis* strains). We did not observe intact, functional partial arrays (e.g., stable one- or two-copy arrays); in contrast, partial arrays were uniformly pseudogenized.

### Association with antiviral, decapping and mitochondrial genes

Across species, STORM insertions consistently occur adjacent to host genes implicated in RNA surveillance, antiviral defense, and mitochondrial function, suggesting that STORM occupies a functionally non-random genomic neighborhood. In the *titani*/*stipitis* clade, STORM occurs between the antiviral helicase SKI2 and the major cytoplasmic decapping enzyme DCP2, separated by ∼13 kb and ∼5 kb, respectively (Fig. 1). In this clade, the array is also flanked by ATP10 and additional mitochondrial-associated genes (Fig. 1). In *S. insectosa*, the antiviral gene SKI3 occurs adjacent to the array; this array-adjacent copy is pseudogenized, but a second, non-array-adjacent SKI3 copy in this species remains intact. RELAX analyses of intact SKI3 orthologs from STORM-containing wood-digesting *Scheffersomyces* clades detected significant relaxation relative to a reference clade (K = 0.75, LR = 10.16, *P* < 0.001). Relaxation was strongest in the *titani*/*stipitis* clade relative to the *lignosus*/*shehatae* clade (K = 0.14, LR = 175.26, *P* < 0.001). Likewise, SKI2 and DCP2, which flank STORM in the *stipitis*/*titani* clade, also showed strong relaxation relative to the *lignosus*/*shehatae* clade. RELAX inferred near-neutralization for both genes when the *stipitis*/*titani* clade was designated as the test group (SKI2: K < 0.01, LR = 129.44, *P* < 0.001; DCP2: K = 0.07, LR = 33.90, *P* < 0.001).

### Genomic mobility and chimerism of STORM arrays

Capsid-domain phylogenies recovered a monophyletic STORM clade within the broader Serinales/L-A-like capsid group (Fig. 3), but with multiple well-supported topological discordances relative to the *Scheffersomyces* species tree (Figs. 3-4). Notably, *S. stipitis* (NRRL Y-7124) grouped with *S. segobiensis* for TLC1-TLC3, but not TLC4, rather than with the conspecific strain *S. stipitis* (CBS 6054), whereas STORM arrays from *S. xylosifermentans*, *S. shehatae*, and *S. insectosa* clustered <u>within</u> the *titani*/*stipitis* clade rather than with the *lignosus*/*shehatae* clade predicted by the species tree.

**Figure 3.**
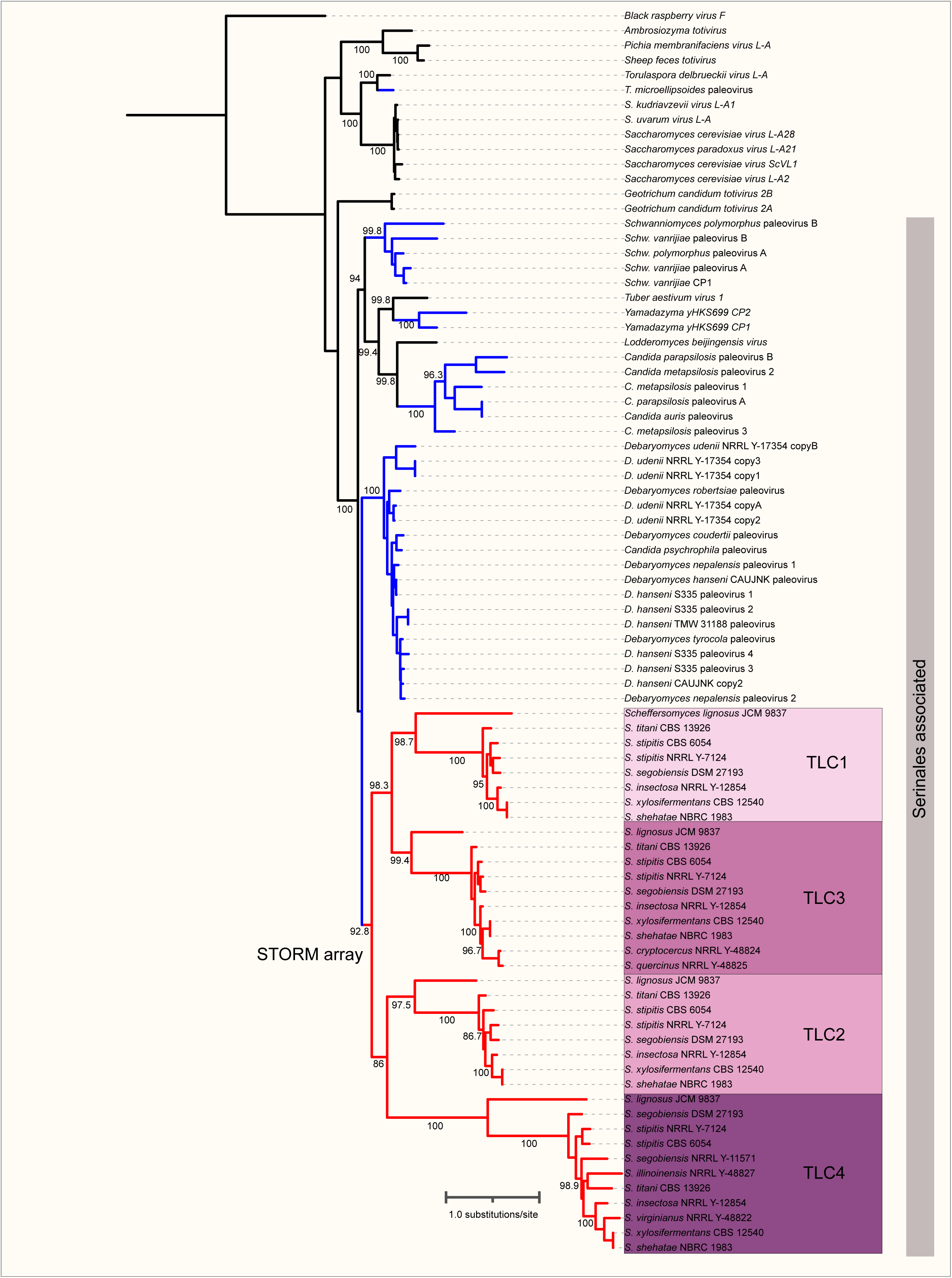
Amino-acid phylogeny of totivirus and totivirus-like capsid proteins. Maximum-likelihood phylogram inferred from amino-acid alignments of capsid domains. Black branches represent capsid sequences from exogenous totiviruses; blue branches correspond to single- or double-copy totivirus-like paleoviruses from non-*Scheffersomyces* yeast genomes; red branches represent the four members of the STORM tandem array (TLC1-TLC4) from species of *Scheffersomyces*. The plum shaded regions highlight the STORM clade and its four capsid-like clades. Tip labels give host species and accession or scaffold identifiers. Node labels show SH-aLRT support (%) for values ≥50. The scale bar denotes 0.5 amino-acid substitutions per site. Sequence metadata for all 84 tips (host species, strain or isolate, primary accession, and re-parsed genomic coordinates) are provided in Supplementary Data S2.

**Figure 4.**
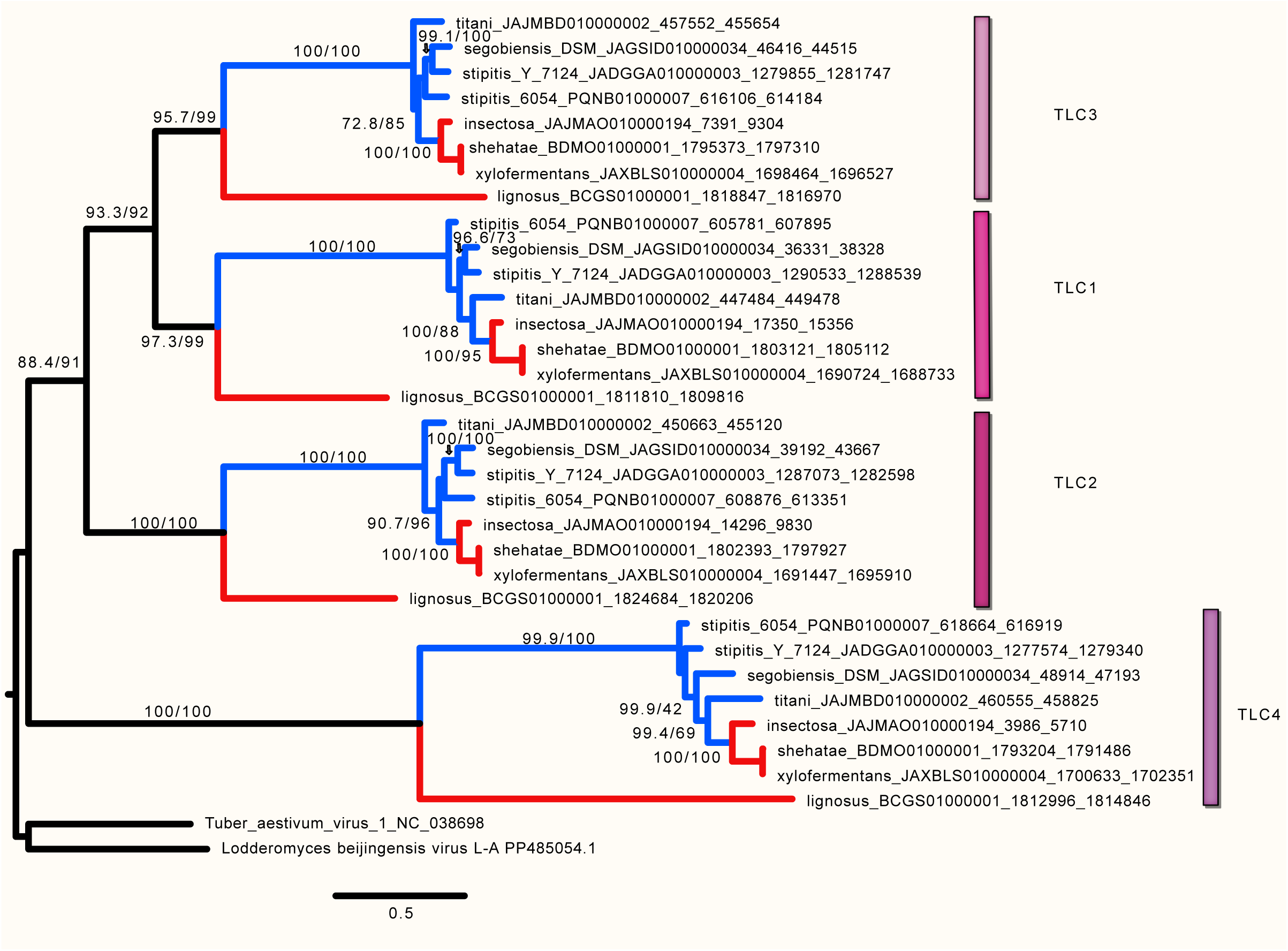
Nucleotide phylogeny of totivirus and totivirus-like capsid genes in Scheffersomyces. Maximum-likelihood tree inferred from nucleotide alignments of capsid and capsid-like paleoviral ORFs. Plum-shaded bars mark the four STORM capsid subclades (TLC1-TLC4) from species of *Scheffersomyces*, with tips labelled by host genome and scaffold coordinates. The tree is rooted with capsid sequences from Tuber aestivum virus 1 and Lodderomyces beijingensis virus L-A (unshaded). Node labels show SH-aLRT / ultrafast bootstrap support (%) for values ≥50, and the scale bar denotes 0.5 nucleotide substitutions per site. The underlying nucleotide alignment is provided as Supplementary Data S4.

Supporting evidence for horizontal transfer includes a pseudogenized ATP10 copy in *S. insectosa* that is more similar to *S. titani/stipitis* clade sequences (where a functional ATP10 flanks STORM; 76% identity to *S. stipitis*) than to the functional ATP10 copy within *S. insectosa* itself (67% identity). The ORF flanking STORM showed highest sequence similarity to Tc1/mariner-like transposases from Mucorales fungi (phmmer: *Rhizopus arrhizus*, *e*-value 1.8 × 10⁻⁹³) and highest structural similarity to a Tc1-like DDE-domain-containing protein from *Lichtheimia ramosa* (foldseek vs. AFDB50, *e*-value 5.68 × 10⁻³²). We designate this transposase family YEET (Yeast Endogenous Element Transposon; Supplementary Figure S1). Phylogenetic analyses recovered two deeply divergent YEET lineages (YEET-α and YEET-β) whose host distributions were discordant with the species tree. Notably, YEET-β elements from *S. insectosa* (including the STORM-adjacent transposase), *S. cryptocercus*, and *S. shehatae* were nested within a clade dominated by *S. stipitis*, *S. segobiensis*, and *S. titani*. These discordant YEET elements showed weak divergence consistent with recent transfers. The terminal inverted repeats (TIRs) of the discordant STORM-associated transposase are of the YEET1a type, which is common in the *segobiensis* clade but not detected in the *lignosus* clade beyond these discordances. Overall, we identified at least five YEET discordances consistent with horizontal transfer among rotten-wood-associated *Scheffersomyces* species (Supplementary Figure S1).

In *S. insectosa*, three YEET TIRs bracket the ATP10 pseudogene and STORM array, consistent with sequential insertion events rather than single integration. Subfamily classification, TIR signatures, and counts are tabulated in Supplementary Data S1 and Supplementary Table S2.

Comparative divergence analyses support YEET-mediated transfer: Kimura 2-parameter distances between *S. insectosa* and *S. segobiensis* were similar for ATP10 (0.310), YEET1a flanking regions (0.285), and YEET1a fragments (0.246), all substantially lower than the ∼0.45 distance for TLC third-codon positions. Within *S. insectosa*, YEET transposases showed recent transpositional activity (mean K2P ≈ 0.053), whereas sister species *S. shehatae* and *S. xylosifermentans* retained identical STORM content (Supplementary Figure S4) but differed markedly in YEET copy numbers and identities, indicating rapid post-speciation loss of transpositional activity. Quantitative comparison with 272 reference nuclear genes (of similar length to TLCs) confirmed the unusual nature of STORM discordances. Only one of 272 reference genes recovered the STORM topology grouping *S. xylosifermentans* with *S. stipitis* (vs. *S. lignosus*) with strong support, and patristic distances between *S. xylosifermentans*(STORM) and *S. lignosus*(STORM) exceeded all reference-gene contrasts (exact Mann-Whitney *P* = 1.53 × 10⁻¹⁰; Fig. 5A).

**Figure 5.**
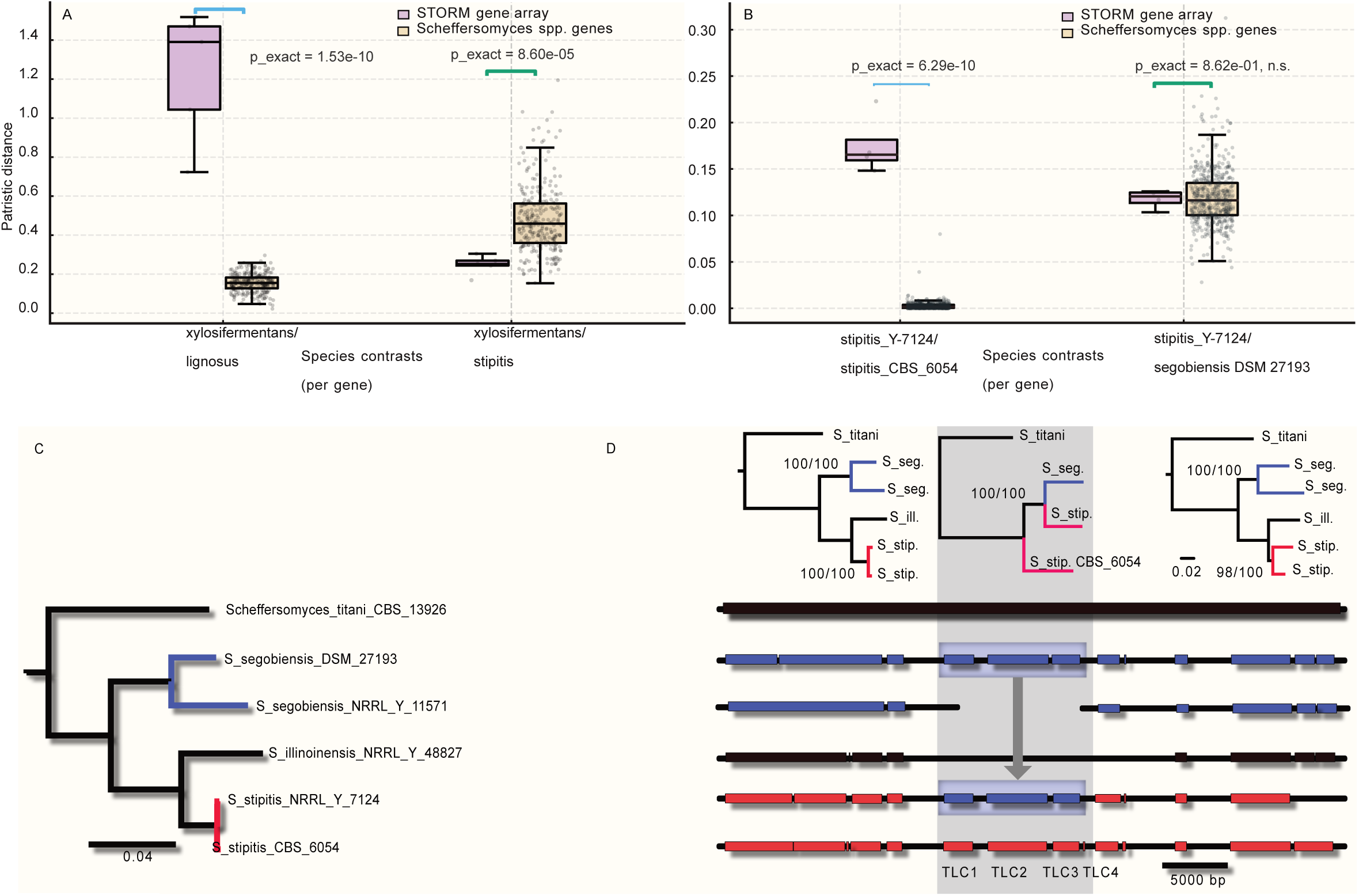
Reticulate transfer of the STORM array inferred from patristic distances and flanking-region phylogenies. (A) Patristic distances between *S. xylosifermentans* and *S. lignosus* (left) and between *S. xylosifermentans* and *S. stipitis* CBS 6054 (right), comparing the STORM array (purple) with 272 reference nuclear genes from *Scheffersomyces* (beige). (B) Patristic distances between *S. stipitis* NRRL Y-7124 and *S. stipitis* CBS 6054 (left) and between *S. stipitis* NRRL Y-7124 and *S. segobiensis* DSM 27193 (right). (C) Species tree for the *titani/stipitis/segobiensis* clade based on concatenated nuclear genes, showing that the two *S. stipitis* strains (red) form a monophyletic group, with monophyletic *S. segobiensis* (blue) more distantly related. (D) Phylogenies of genomic regions flanking the STORM array (upper panels) and corresponding BLASTn similarity blocks using the *S. titani* region (JAJMBD010000002.1:430000-519758) as a query (lower panel). Trees from upstream and downstream segments recover the expected species relationships, with *S. stipitis* strains forming a clade and *S. segobiensis* outside (blue vs red), whereas the array region itself (gray block) is discordant. The gray arrow represents the proposed direction of gene transfer.

Similarly, for *S. stipitis* strains, only one of 523 reference genes showed the STORM-like discordance, whereas TLC1-TLC3 divergence between conspecific *S. stipitis* strains exceeded all reference-gene distances (Fig. 5B). The species tree for the *titani*/*stipitis*/*segobiensis* clade places the two *S. stipitis* strains as monophyletic sister taxa with monophyletic *S. segobiensis* as outgroup (Fig. 5C). Trees from regions upstream and downstream of TLC1-3 recover this expected species topology with strong bootstrap support (Fig. 5D), but the tree from TLC1-3 itself places *S. stipitis* NRRL Y-7124 with *S. segobiensis* rather than with conspecific *S. stipitis* CBS 6054. There is a sharp topological transition at the TLC1-3 boundaries, consistent with a localized transfer event that replaced the STORM region of NRRL Y-7124 while leaving the surrounding genomic context intact. This partial-replacement pattern, with the TLC3-TLC4 junction marking an apparent internal breakpoint, is consistent with recombination between donor and recipient arrays during the transfer event.

### Tandem duplication generates asymmetric evolution

We next compared relative amino-acid divergence among (i) extant L-A-like totiviruses, (ii) singleton totivirus-like paleoviruses outside STORM, and (iii) the four STORM paralogs. Using maximum-likelihood capsid trees (related totivirus sequences provided a root; see Methods), root-to-tip distances were greatest for TLC4 (mean 5.05 expected amino-acid substitutions per site; Fig. 6A). TLC2 and TLC3 showed intermediate distances (means ∼3.8-3.9), and TLC1 was lowest within STORM (mean 3.66). Non-STORM paleoviruses were the least divergent group overall. L-A-like totiviruses were more divergent than non-STORM paleoviruses but remained markedly less divergent than STORM paralogs (Fig. 6A). Only one pairwise contrast lacked significance, TLC2 with TLC3 (pairwise Mann-Whitney tests with FDR correction;

**Figure 6.**
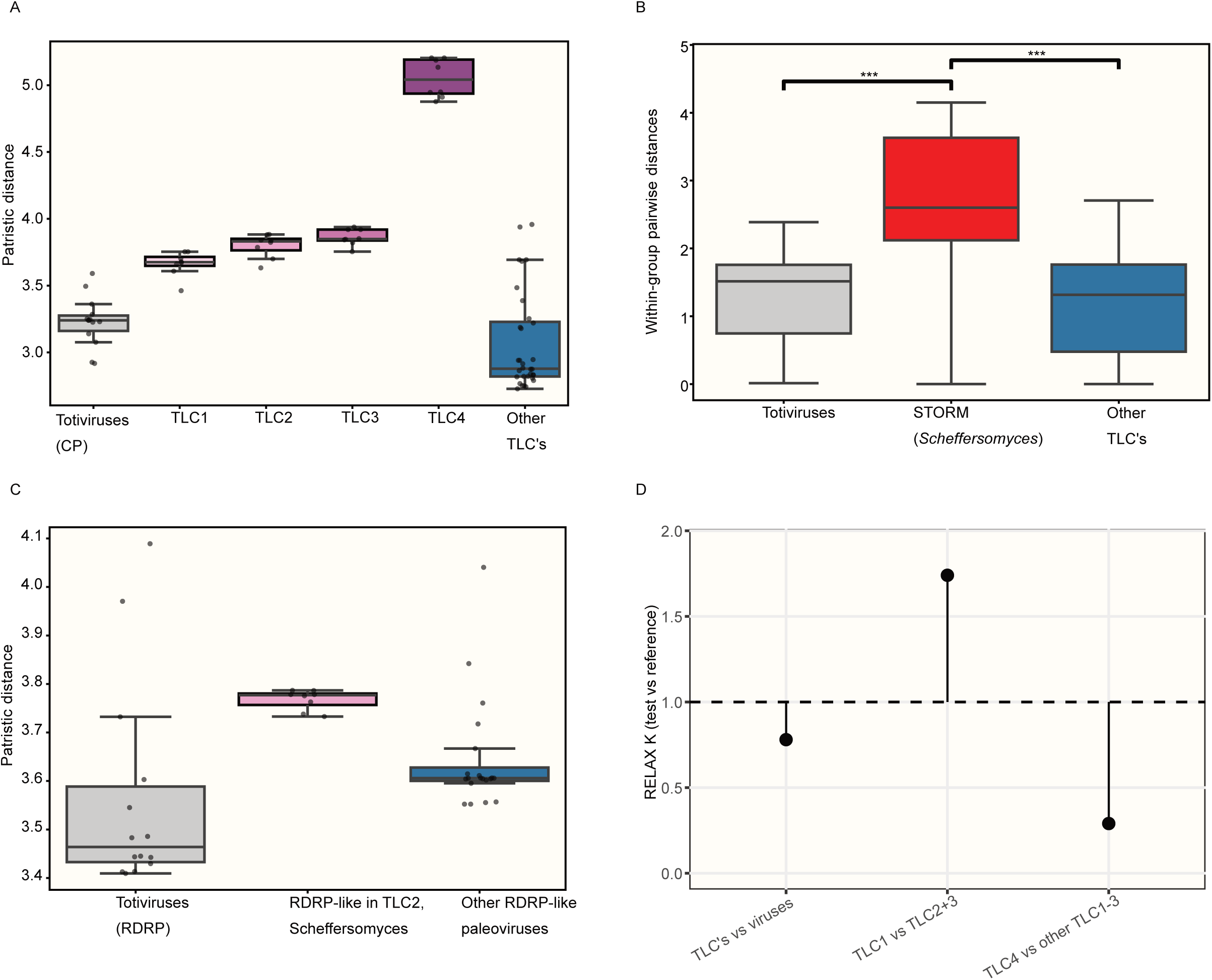
Multimodal evolutionary rates and selection on STORM capsids and RdRp. (A) Root-to-tip patristic distances for capsid (CP) amino-acid trees, showing exogenous totiviruses (gray), the four STORM copies TLC1-TLC4 (plum), and other totivirus-like capsid paleoviruses (blue). (B) Within-group pairwise patristic distances for capsid sequences, comparing exogenous totiviruses (gray), STORM capsids from *Scheffersomyces* (red), and other capsid paleoviruses (blue). (C) Root-to-tip patristic distances for RdRp trees, comparing exogenous totiviruses (gray), the RdRp-like paleovirus from *Scheffersomyces* STORM (plum), and other RdRp-like paleoviruses (blue). (D) RELAX tests of selection intensity (K) for capsid genes. From left to right: all STORM TLCs vs exogenous viruses; TLC1 vs TLC2+TLC3; and TLC4 vs other TLC1-3. K < 1 indicates relaxation in the test set; K > 1 indicates intensification. The dashed line marks K = 1.

Supplementary Table S6). Filtering the alignment with ClipKIT (keeping constant sites, kpic smart gap) removed 25.4% of sites and marginally reduced the patristic distances but retained the relative divergence patterns (Supplementary Figure S2). Within-group distance comparisons further showed that STORM paralogs are significantly more divergent than both extant viruses and non-STORM paleoviruses (Fig. 6B). Anchoring these distances to host divergence times highlights the unusual tempo of STORM evolution. The deep ingroup common ancestor of the sampled totivirus-bearing yeasts is estimated at ∼225 MYA, whereas the *Scheffersomyces* clade common ancestor is ∼54 MYA (Kumar et al. 2022). Despite this substantially shorter host timescale, STORM accumulated greater amino-acid divergence than the sampled exogenous totivirus lineages, with TLC4 alone exceeding the mean root-to-tip divergence of any exogenous totivirus in our sampling. A similar pattern was observed for RdRp domains (which, for STORM, are only present in TLC2): STORM-associated RdRp sequences exhibited greater patristic distances than related viral and paleoviral RdRps (Fig. 6C). Together, these results indicate that tandem duplication generated an array in which paralogs occupy distinct positions along a long-term evolutionary-rate spectrum, with TLC4 exceeding related viral lineages in apparent amino-acid divergence.

### Relaxed selection and structural evolution of an accelerated copy

To test whether elevated TLC4 divergence reflects relaxed purifying selection rather than pervasive positive selection, we applied codon-based tests for selection and relaxation (Fig. 6D). RELAX analyses comparing STORM capsid branches to extant viral capsid branches detected a non-significant trend toward relaxation (K = 0.78, LR = 3.16, P = 0.075).

Focusing on individual array members, TLC4 branches showed strong evidence of relaxation relative to the remaining TLC branches (K = 0.29, LR = 74.12, *P* < 0.001). In contrast, TLC1 branches showed significant intensification relative to TLC2 and TLC3 (K = 1.74, LR = 16.47, *P* < 0.001). Under the RELAX alternative models, the inferred purifying class on TLC1 branches had substantially lower dN/dS (∼0.03-0.04) than TLC2 and TLC3 (∼0.10-0.13). In contrast, TLC4 showed a shift toward substantially higher dN/dS (∼0.5) together with an increased fraction of sites assigned to a high-ω class. Using Contrast-FEL across 658 codons, we found extensive site-specific differences between TLC4 and the rest of the array: TLC1 vs TLC4 differed at 84 sites (24 FDR-significant at q ≤ 0.2), and TLC2/3 vs TLC4 differed at 71 sites (28 FDR-significant). By contrast, TLC1 vs TLC2/3 yielded no FDR-significant differences (44 nominal sites at *P* ≤ 0.05, but q > 0.2). Together, these results support a distinct relaxed selective regime on TLC4, whereas TLC1 and TLC2/3 share broadly similar purifying selection, with significant intensification on TLC1.

To assess whether TLC4 divergence compromises protein structure, we predicted structures for each capsid-like paralog with AlphaFold3 and compared them to each other and to known totivirus capsids using Foldseek (Abramson, et al. 2024; van Kempen, et al. 2024). TLC1-TLC3 retained a conserved totivirus-like capsid architecture, including the core β-sandwich and surrounding α-helices. However, these copies lacked residues associated with the decapping loop and showed corresponding alterations in loop structure (Fig. 7A,B). TLC4 preserved key residues and the overall decapping-loop configuration but showed the largest deviations in surface loops and protruding domains. Although TLC4 uniquely lacked the conserved totiviral α1- α2 (helix-linker-helix) motif (Fig. 8B) and had significantly lower TM scores than TLC1 (Fig. 8A), its TM score still indicates strong structural conservation relative to exogenous viral capsids. Thus, despite pronounced sequence divergence, TLC4 differs structurally from other STORM copies primarily through loss of the conserved helix-linker-helix motif while retaining the decapping loop.

**Figure 7.**
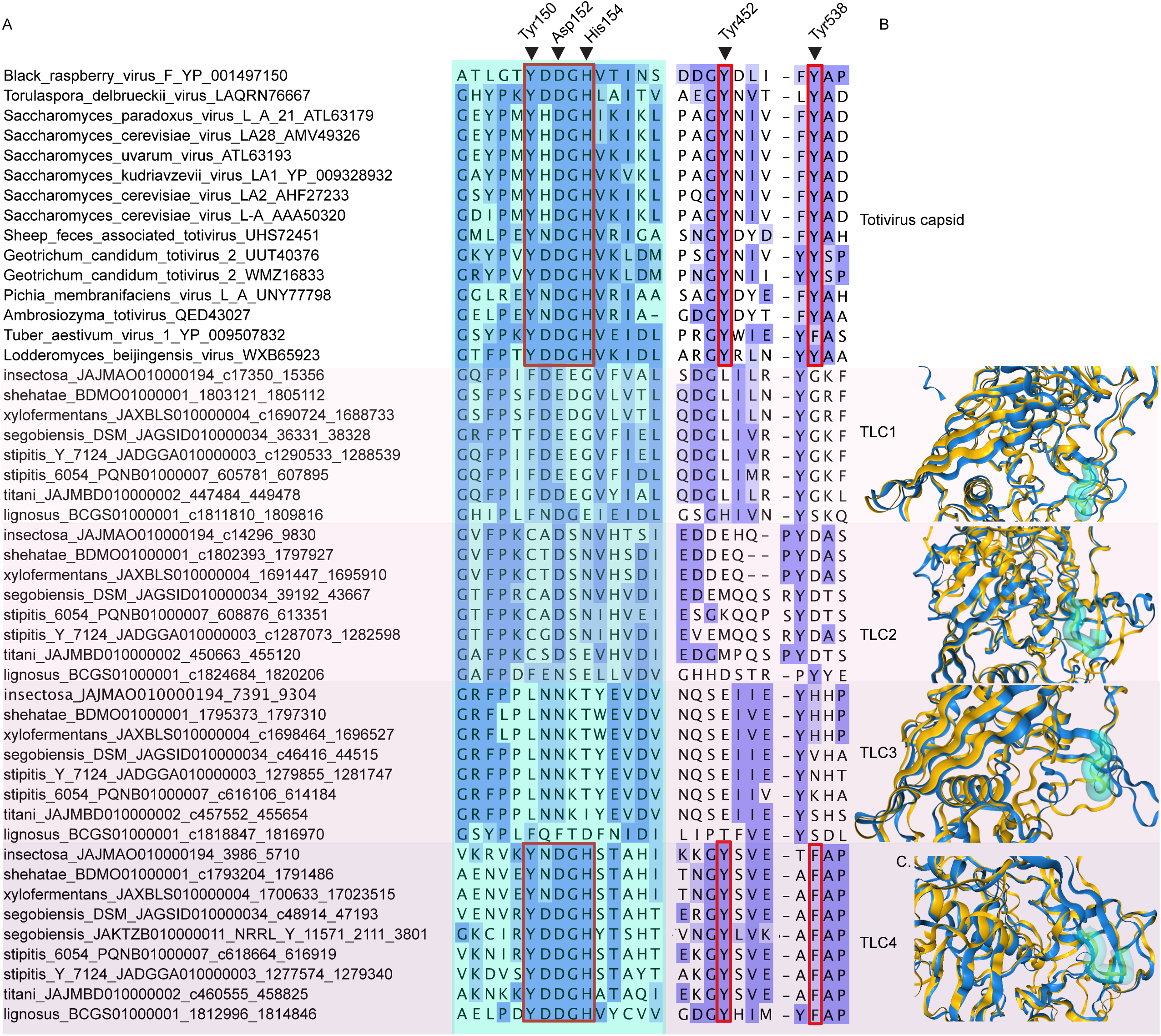
Conservation of the decapping-loop motif and local capsid structure in STORM paleoviruses. (A) Amino-acid alignment of the region encompassing the putative decapping loop in totivirus capsid proteins and totivirus-like capsids from *Scheffersomyces* and other yeasts. Viral capsids are shown at the top, followed by single-copy paleoviruses, and then the four STORM copies (TLC1-TLC4; shaded background). Conserved residues implicated in decapping activity are highlighted by red boxes. (B) Local 3D structures of the capsid region containing the presumptive decapping loop. The loop is highlighted as a teal tube. For comparison, the predicted structure of the Lodderomyces beijingensis virus capsid is shown in yellow, and the AlphaFold3-predicted capsid structure from a representative STORM array (*S. titani*) is shown in blue. TLC4 alone retains both the conserved decapping-loop motif and the loop in its canonical structural position.

**Figure 8.**
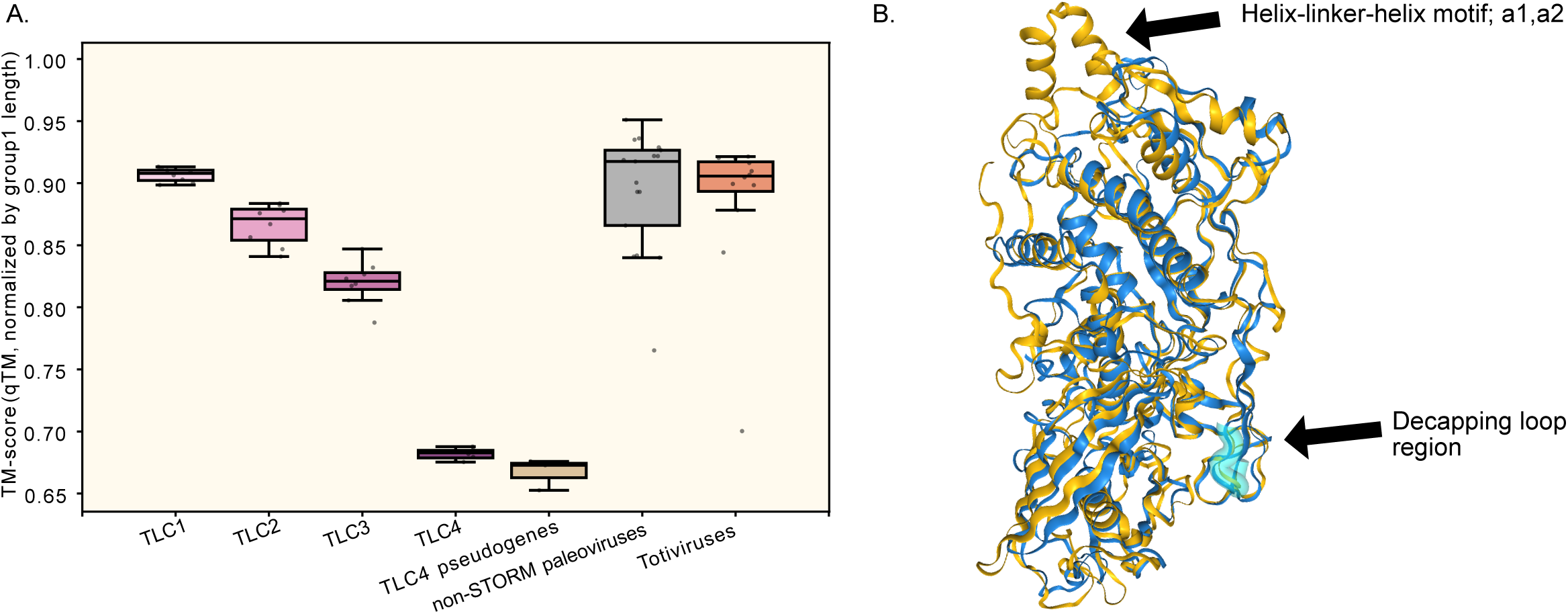
Structural similarity of STORM capsids to viral capsids and loss of the helix-linker-helix motif in TLC4. (A) TM-scores (qtmscore, normalized by viral query length) between AlphaFold paleoviral capsid predictions and exogenous totivirus references. Each paleoviral structure was assigned the highest TM-score obtained against any Serinales-associated viral reference. Non-STORM TLCs were filtered to retain only high-coverage alignments (query coverage ≥80%; 32 → 17 structures). Group differences were significant overall (Kruskal-Wallis, P < 0.001), with TLC4 ORFs showing significantly lower structural similarity to viral capsids than TLC1 and non-STORM paleoviruses (pairwise Mann-Whitney tests; see Supplementary Data S7). (B) Representative structural overlay of TLC4 (*S. titani*; blue) with a totivirus capsid (*Lodderomyces beijingensis virus*) template (yellow).

### Condition-dependent expression of STORM and neighboring host genes

To determine whether STORM is transcriptionally active, we performed targeted RNA-seq quantification across five *Scheffersomyces* species and strains under varying growth conditions (see Methods). Within each sample, TPM values were normalized to ACT1 to facilitate cross-species comparisons (Fig. 9). Across taxa and conditions, TLC1 was consistently the dominant STORM transcript (typically 20-35% of ACT1; range ∼9-35%). TLC3 averaged ∼4% of ACT1, whereas TLC2 and TLC4 were generally lower (typically 0.4-3% of ACT1), with the notable exception of TLC4 in *S. insectosa*. TLC4 expression was highest in *S. insectosa* (∼11% of ACT1), although transcriptomes for this species lacked biological replication, suggesting a possible effect of local genomic context. ACT1 remained stable across conditions and species, supporting its use as a reference, whereas TEF1 was condition-sensitive in *S. stipitis* but relatively stable in *S. xylosifermentans*.

**Figure 9.**
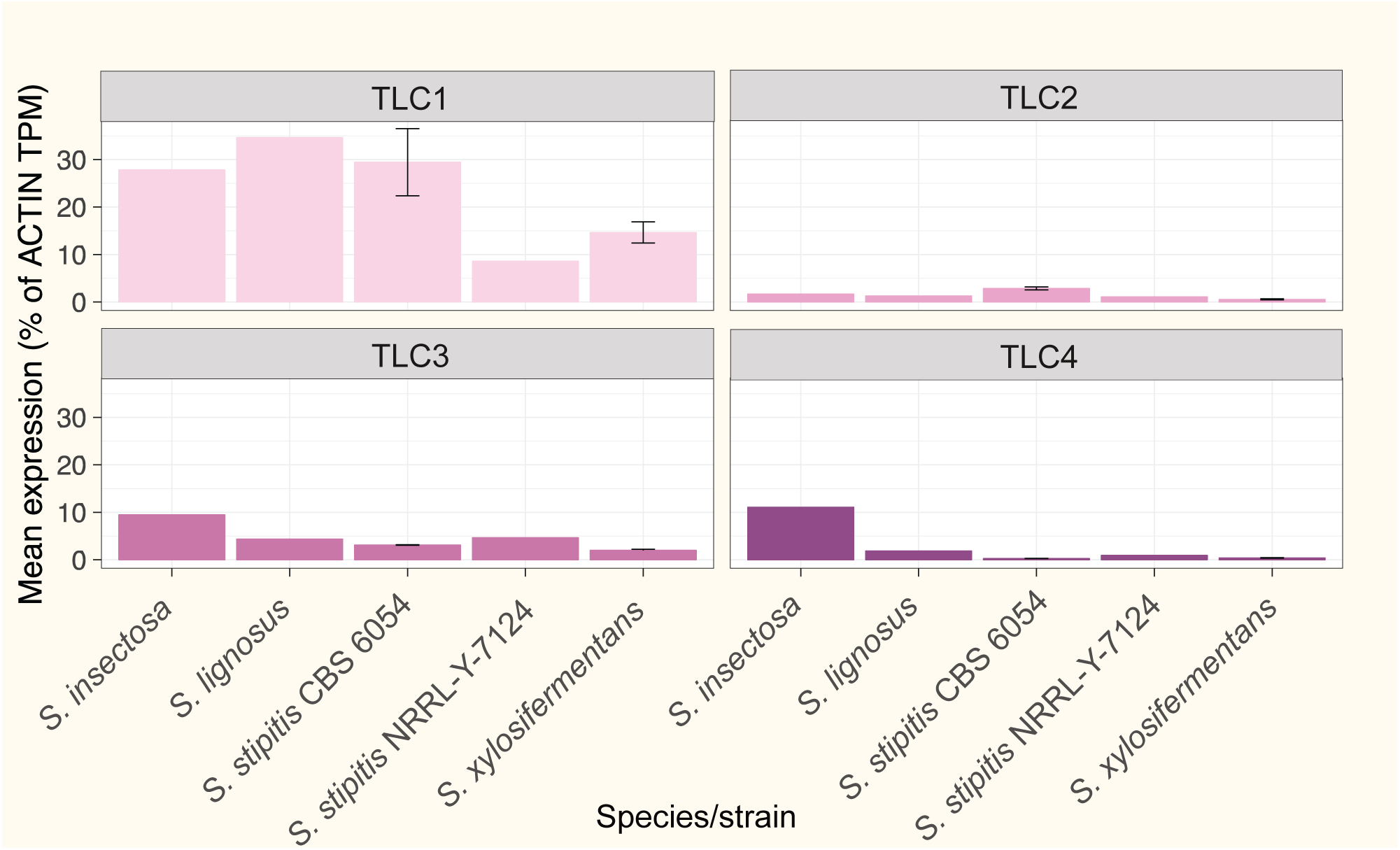
Cross-species expression of STORM capsid genes in *Scheffersomyces*. Bar plots show mean expression of each STORM member (TLC1-TLC4; separate panels) across five *Scheffersomyces* species and strains, expressed as a percentage of *ACT1* (ACTIN) transcript abundance (TPM; y-axis). Error bars indicate ± SE where biological replicates were available.

In *S. xylosifermentans*, TLC1 and TLC4 were ∼2-fold higher under baffle conditions (higher O₂ transfer), TLC2 was ∼1.5-fold higher under shake, and TLC3 showed modest baffle bias (Fig. 10A). Among host genes in the same species, SKI2 was induced ∼3-fold under shake, SKI3 decreased modestly, and DCP2 remained stable (Supplementary Figure S3). Several nuclear and mitochondrial genes (SMC5, PEX19, TFC1, PTC5) were also induced under shake, whereas MSE1 and PDH were higher in baffle. Notably, higher TLC1 and TLC4 expression under baffle coincided with higher SKI2 expression under shake - opposing rather than coordinated responses along the same antiviral axis.

**Figure 10.**
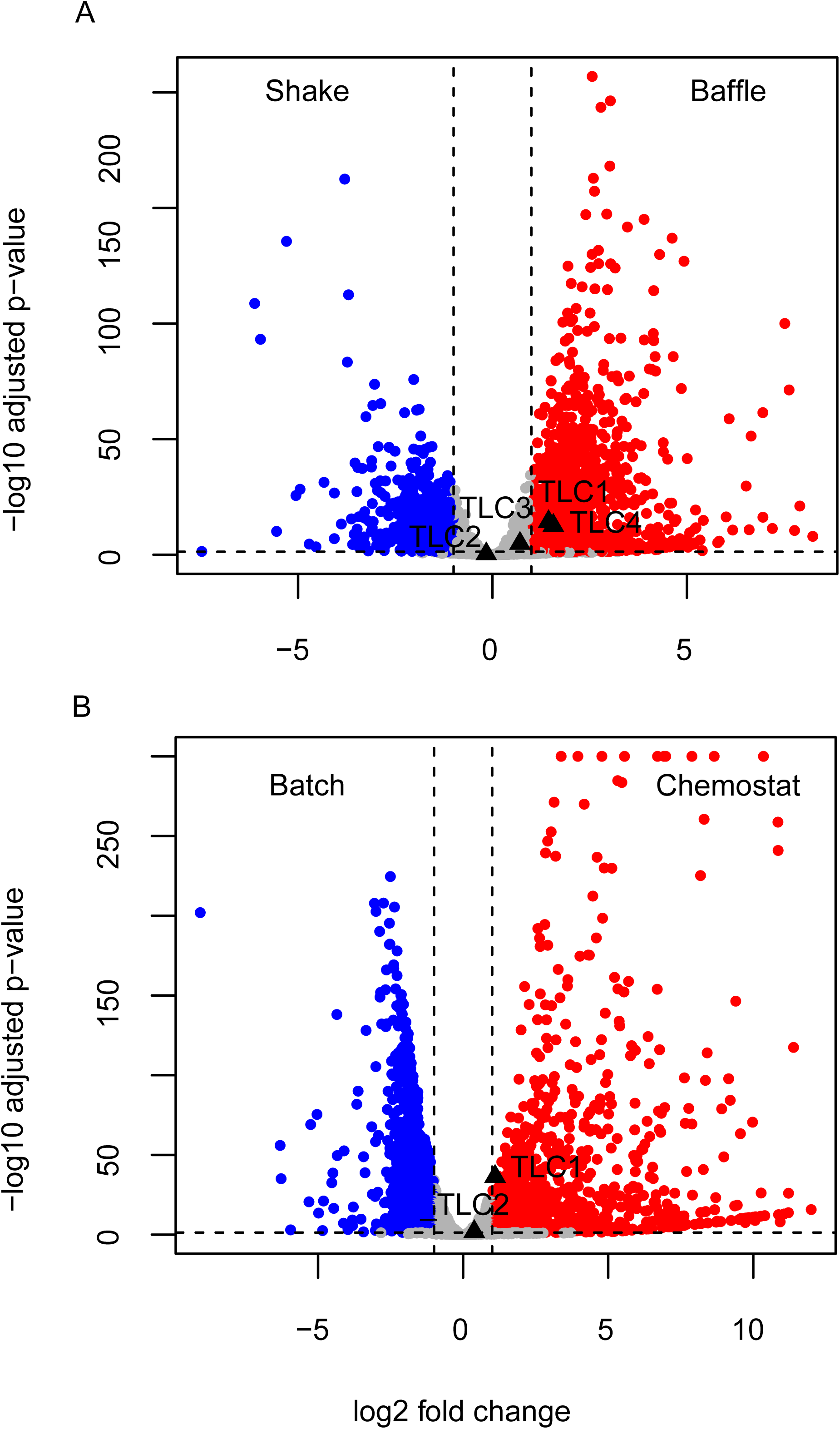
Differential expression of STORM capsids in *Scheffersomyces* across growth regimes. (A) Volcano plot for *S. xylosifermentans* comparing shake vs baffle (higher O₂ transfer) cultures.Each point represents a gene; the x-axis shows log₂ fold change (positive values = higher expression in baffle, negative = higher in shake), and the y-axis shows -log₁₀ adjusted *P*-value. Red and blue points indicate significantly upregulated genes in baffle and shake, respectively (FDR threshold indicated by the horizontal dashed line; vertical dashed lines mark fold-change cutoffs). Triangles highlight the four STORM capsid genes (TLC1-TLC4). (B) Volcano plot for *S. stipitis* CBS 6054 comparing batch-biased (blue) vs chemostat-biased (red) expression, plotted as in (A) (positive log₂ fold change = higher expression in chemostat).

In *S. stipitis* (CBS 6054), TLC1 and TLC2 were significantly upregulated in chemostat relative to batch (∼2.6-fold and ∼1.6-fold respectively), whereas TLC3 and TLC4 changed little (Fig. 10B). In the same species, SKI3 and DCP2 were significantly higher in batch, whereas SKI2 showed a weak, non-significant decrease in chemostat (Supplementary Figure S3). Thus, elevated TLC1 and TLC2 in chemostat coincided with reduced SKI3 and DCP2 - a pattern opposite to that observed in *S. xylosifermentans*. Together, these results show that expression is strongly condition dependent and that the same array can exhibit different regulatory partitioning across species: in *S. xylosifermentans* both ends of the array respond to mixing regime, whereas in *S. stipitis* expression is front-loaded toward the 5′ portion of the array under chemostat. Rather than acting through a single conserved antiviral pathway, STORM appears to participate in distinct regulatory contexts across *Scheffersomyces* lineages, with functional associations varying by species and growth condition.

## Discussion

### Tandem paleoviral arrays as evolutionary modules

Our results reveal that paleoviruses derived from nonretroviral RNA viruses can follow unexpected evolutionary trajectories, including asymmetric tandem array evolution in which gene copies accumulate greater amino-acid divergence than their sampled viral relatives while preserving ancestral capsid-like structures. STORM represents a long-lived virus-derived module that has persisted for roughly 15 million years in the syntenic *S. titani* clade (and more than 50 MY if *S. lignosus* is included), remained horizontally mobile among yeast genomes, and become transcriptionally integrated into host regulatory networks. Because the array encodes both polyprotein-like and capsid-like proteins, STORM may retain some of the proposed stress tolerance benefits associated with L-A viruses without the chronic maintenance costs and proteotoxicity imposed by exogenous totiviral infections. Tandem array formation in yeast is more broadly associated with increased stress tolerance (Hsieh, et al. 2025), suggesting that STORM’s architecture may itself confer adaptive benefits independent of its viral origin. STORM therefore represents a domesticated viral module whose internal paralogs have diverged under distinct selective regimes.

Time-dependent rate models such as the Prisoner-of-War (PoW) framework predict that, over long timescales, RNA virus substitution rates decay toward the host background as constraints and saturation proceed. Viral capsids remain under strong structural (and sequence) constraints associated with virion assembly and genome packaging (Luque, et al. 2018). Accordingly, long-persisting functional paleoviral genes should ultimately approach host-like evolutionary rates regardless of PoW dynamics. The exogenous totiviruses sampled here span deep phylogenetic timescales and may themselves represent long-term PoW-decayed rates rather than true short-term viral rates. Non-STORM TLC-like paleoviruses evolve at rates similar to, or only slightly slower than, this exogenous totivirus reference (Fig. 6A), a pattern that is observationally consistent with both reference and paleoviruses having settled into a long-term equilibrium near, but not necessarily at, host rates. STORM is a clear exception against this baseline, with TLC4 evolving fastest of all (Fig. 6A). Because integrated paleoviral genes are constrained to evolve no faster than the host’s neutral mutation rate, TLC4’s elevation above the totivirus reference implies that purifying constraint has been substantially relaxed (consistent with the K < 1 signal from RELAX), letting accumulated divergence climb toward the neutral ceiling rather than remaining at the more constrained equilibrium reached by other paleoviruses. Tandem duplication of domesticated viral genes creates extra copies that can explore sequence space under relaxed constraints without jeopardizing core function.

Despite accelerated evolution and relaxed selection, the most divergent member (TLC4) retains a recognizable totivirus-like capsid fold and key decapping-related motifs (Fig. 7). This pattern resolves a paradox. TLC4 is simultaneously the most divergent member of the array and the only paralog retaining the ancestral decapping-loop motif (lost from TLC1, TLC2, and TLC3). The selective retention of predicted decapping loop residues in TLC4, despite their loss from the more sequence-constrained paralogs (TLC1-TLC3), argues against simple degenerative drift and instead suggests copy-specific constraint on this feature. One possibility is that decapping activity confers a fitness benefit through modulation of host mRNA turnover even in the absence of active viral infection. That TLC4 is consistently retained after movement into distinct genomic contexts further supports its functional importance within the array. This pattern is expected from classic models of duplicate evolution in which one copy maintains ancestral function while another explores novel sequence space (Ohno 1970; Zhang 2003; Innan and Kondrashov 2010). In STORM, however, this asymmetry is reversed from the naive expectation: the most divergent paralog (TLC4) is the one that retains the ancestral decapping motif, whereas the more sequence-constrained paralogs (TLC1-TLC3) have lost it. The Contrast-FEL site-level test reinforces this two-class pattern. TLC4 differs from the rest of the array at dozens of FDR-significant codons, whereas TLC1 and TLC2/3 are statistically indistinguishable from each other, indicating that the elevated rate on TLC4 reflects relaxed constraint across much of the protein rather than uniform decay.

### STORM, RNA-decapping pathways, and potential host benefits

Compared with hosts that possess adaptive immune systems and rapidly evolving antibodies (Tenthorey, et al. 2022), yeasts seem poorly equipped to engage in classic arms races with RNA viruses. The evolution of their antiviral genes and innate restriction factors is constrained by the comparatively modest evolutionary rates of eukaryotic genomes. Nevertheless, expression of TLC-like paleoviral genes can interfere with replication of the L-A virus (Warner, et al. 2018). A module of differentiated tandem antiviral genes such as STORM could expand this repertoire, preserving multiple antiviral solutions and permitting the rapid exploration of new ones under relaxed selection.

The repeated association of STORM with antiviral and decapping genes (e.g. SKI2, SKI3, DCP2), together with conserved decapping-related motifs in TLC4, suggests a functional link between the array and host antivirus/RNA-decay machinery. SKI2, SKI3, and DCP2 show strong relaxation in the *stipitis*/*titani* clade, whereas TLC4 retains predicted decapping activity. This pattern is consistent with partial attenuation or remodeling of host RNA-decay pathways in STORM-containing lineages, with selective preservation of viral-cap-processing features.

STORM may therefore act not simply by competing with SKI/XRN1, but by augmenting or regulating these pathways to throttle viral replication. Differences in condition-dependent expression of STORM and antiviral genes between *S. xylosifermentans* and *S. stipitis* suggest that each lineage has tuned this module differently. STORM proteins could therefore modulate viral replication through lineage-specific interactions with viral RNAs, viral proteins, or host stress-response pathways.

Expression analyses show that STORM is not a silent genomic relic. Across *Scheffersomyces*, TLC1 reaches moderate to high expression levels (often tens of percent of ACT1), with proteomic evidence confirming translation in *S. stipitis* (Taylor et al. 2013). TLC3-TLC4 are lower but consistently expressed and condition-responsive.

Several features distinguish TLC4 from a pseudogene under neutral decay: STORM TLCs retain intact ORFs while neighboring co-transferred genes (ATP10, transposases) are pseudogenized; recent TLC4 pseudogenes are structurally more divergent than the intact TLC4 copies; and intact copies remain transcriptionally active. Despite extreme primary-sequence divergence, TLC4 maintains a conserved totivirus-like capsid fold, including the predicted decapping loop region, with high structural similarity to exogenous capsids and minimal variation across deeply divergent STORM contexts. Together, these observations indicate that TLC4 is not a decaying pseudogene, but a rapidly evolving STORM component whose coding sequence and capsid-like structure remain under selective constraint despite extreme sequence divergence.

The coordination of STORM expression with nuclear maintenance, mitochondrial, and antiviral pathways suggests the array functions as part of a broader stress-response network rather than a dedicated antiviral module. Given that DCP2 in yeast also functions to repress a large cohort of OXPHOS transcripts and suppresses respiratory metabolism (Vijjamarri, et al. 2023), the physical proximity of the STORM array to DCP2 suggests potential integration into host metabolic regulatory circuits. Taken together, the STORM-SKI2-DCP2 neighborhood may represent a functional RNA surveillance and metabolic coordination domain rather than a coincidental genomic arrangement.

### Reticulate evolution and losses of the STORM array

Our species tree agrees with Opulente et al. (2024), but phylogenetic discordances and the mosaic structure of STORM indicate reticulate evolution. Several *Scheffersomyces* species co-occur in insect guts and decaying wood (Urbina, et al. 2013; Ceja-Navarro, et al. 2014), contexts conducive to interspecific exchange. Two mechanisms could explain localized reticulation: hybridization followed by loss of heterozygosity (LOH), or horizontal transposon-mediated transfer (HTT).

Two features of fungal biology converge on the transposon hypothesis. The YEET transposase belongs to the Tc1/*mariner* superfamily, which Romeijn et al. (2026) identified as the single most pervasively horizontally transferred TE superfamily across fungi, despite retroelements being more abundant overall; and HTTs are overrepresented between closely related taxa (Romeijn, et al. 2026). The combination of an HTT-prone vehicle and shared microhabitats among closely related congeners offers a parsimonious explanation for why we find this signature in *Scheffersomyces* even though it is rare in the broader Saccharomycotina. Multiple lines of locus-level evidence support HTT with a paleoviral cargo. The STORM array in *S. insectosa* is bracketed by YEET transposable elements that also flank a pseudogenized *ATP10* copy more similar to *S. titani*-clade orthologs than to the conspecific functional copy. The concordant K2P distances among ATP10, YEET1a flanking regions, and YEET1a fragments (Fig. 5) are consistent with a shared transposition event. Cross-clade YEET-β placements involving *S. insectosa*, *S. cryptocercus*, and *S. shehatae* indicate element mobility across host lineages. Additionally, in sister species *S. shehatae* and *S. xylosifermentans*, STORM content is identical, but flanking YEET elements have diverged in copy number and sequence, consistent with post-transfer silencing. TE-mediated transfer is more parsimonious than hybridization-LOH: our 271/272 and 522/523 reference-locus scans recover no additional introgressed blocks at the size expected under hybridization-LOH, though we cannot exclude single-gene introgressions below the resolution of our gene-tree comparisons.

Incomplete lineage sorting (ILS) is unlikely to explain the STORM discordance on both empirical and mechanistic grounds. Empirically, the multispecies coalescent predicts discordance frequencies determined by ancestral *N*ₑ and internode length, yet genome-wide discordance is rare for the relevant nodes: 271 of 272 reference loci between *S. xylosifermentans* and *S. lignosus*, and 522 of 523 between the two *S. stipitis* strains, recover the species tree. Under parameterizations consistent with this background, the probability of repeatedly recovering the STORM-like topology across multiple tandem paralogs is extremely low. Mechanistically, ILS assumes vertical inheritance at orthologous loci, whereas STORM occurs in multiple genomic contexts across our sampling (including different chromosomes in the two *S. stipitis* strains). Together, these observations point to a non-coalescent explanation, consistent with mobility of the array, potentially via transposon-mediated transfer, as supported by flanking signatures and convergent K2P distances of co-located *ATP10* and *YEET1a*.

Despite earlier reports of poorly supported non-monophyly for S. stipitis CBS 6054 capsid paleoviruses (Taylor and Bruenn 2009; Brejová et al. 2024), our expanded taxon sampling and updated phylogenetic analyses recover a monophyletic STORM clade (Fig. 3).

The breakpoint-equivalent evidence from windowed phylogenies and paralog-by-paralog resolution of the chimeric S. *stipitis* NRRL Y-7124 array (Figs. 4 and 5) is consistent with intrarray recombination at the TLC3-TLC4 junction during transfer. Gene conversion among TLC paralogs within a single array is unlikely to contribute substantially beyond these transfer-coincident events, because the >50% protein divergence between TLC4 and the remaining paralogs exceeds the sequence similarity typically associated with efficient gene conversion, and the persistent extreme divergence of TLC4 argues against ongoing homogenizing forces.

Tandem paralog arrays provide abundant substrate for homologous exchange, and secondary reshuffling could decouple the evolutionary histories of individual TLC copies even if the array initially moved as a unit. We therefore cannot exclude the possibility that some STORM topologies reflect chimeric evolutionary histories superimposed on broader patterns of transposon-associated mobility. Nevertheless, the shared monophyly of the array, conserved tandem architecture, and concordant flanking signatures together support movement of STORM as a mobile module rather than independent origins of individual TLC genes.

STORM shows recurrent losses across lineages, indicating context-dependent costs. Pseudogenes in the *S. cryptocercus*/*S. quercinus*/*S. virginianus* group and losses in *S. illinoinensis* and *S. segobiensis* represent multiple independent events. For *S. segobiensis*, STORM loss coincides with a putative killer system (Taylor, et al. 2013), suggesting that domesticated capsid functions could become incompatible with exogenous viral elements in some genomic backgrounds.

### Paleoviruses as experimental systems for time-dependent rate dynamics

STORM demonstrates that nonretroviral RNA paleoviruses can follow evolutionary trajectories distinct from those of exogenous viruses after host integration and tandem duplication.

Its viral progenitors are structurally and functionally well characterized; the array exists in multiple copies, species, and genomic contexts; and its minimum age can be constrained by host divergence times. Together, these features allow comparative tests of how duplication, genomic context, and regulatory integration mediate the transition from virus-like to host-like evolution. Our findings add complexity to current models of time-dependent viral evolution and virus-host coevolution: that active host genes derived from RNA viruses and subject to relaxed or shifted selection after duplication may evolve more rapidly at the protein level than strongly constrained viral genes. We expect similar domesticated modules to exist for other nonretroviral RNA viruses, suggesting that relaxed host-associated evolutionary regimes for virus-derived paralogs may represent an underappreciated component of virus-host coevolution.

## Materials and Methods

### Data collection and identification of totivirus-like sequences

We compiled totivirus and totivirus-like sequences from the NCBI nonredundant (nr) protein database using BLASTP with yeast L-A capsid and RdRp proteins as queries (capsid: AAA50320.1; RdRp: AAA50321.1) and additional related viral sequences. Candidate exogenous totiviruses were retained if they contained (i) a recognizable totivirus-like capsid region and (ii) conserved RdRp motifs. We use the term “paleovirus” rather than “endogenous viral element” because “endogenous virus” has a distinct meaning in the totivirus literature, referring to persistent vertically transmitted dsRNA viruses (e.g., Fujimura and Wickner 1989; Schmidt et al. 2024; Hsiao et al. 2025). We use NIRV (non-retroviral integrated RNA virus; Taylor and Bruenn 2009) synonymously in figure labels. To identify totivirus-like paleoviruses in yeast genomes, we screened publicly available assemblies from CTG-clade yeasts (Debaryomycetaceae; Serinales) using TBLASTN with totivirus capsid and RdRp queries. We retained hits with E-value < 1×10⁻⁵ and alignment length ≥ 300 amino acids. Candidate loci were refined by ORF prediction (minimum ORF length 300 nt) under the appropriate yeast nuclear genetic code (i.e., genetic code 12 for Serinales).

### ORF calling and array annotation

Open reading frames were identified using NCBI ORFfinder (minimum ORF length 300 nt, genetic code 12, nested ORFs ignored). Capsid-like ORFs were identified based on similarity to totivirus capsid proteins and the presence of conserved capsid-associated motifs. A polyprotein was defined as an ORF containing both capsid-like and RdRp-like domains. Annotated coding sequences were exported in FASTA format for downstream analyses.

### Microsynteny and macrosynteny analysis

To assess microsynteny around STORM arrays, we extracted STORM-containing genomic regions (e.g., *S. titani* strain CBS 13926, JAJMBD010000002.1:430000-519758) and searched against *Scheffersomyces* genome assemblies using BLASTN. Flanking ORFs were predicted as described above.

Contiguous high-scoring blocks (combined bit score ≥ 200, E < 1e−5) were used to delineate syntenic regions. Genomes with syntenic flanks but lacking array sequences were scored as array-lacking, whereas those with both flanks and internal capsid-like hits were scored as array-present. Putative flanking genes were annotated using ORF prediction and BLASTP searches against UniProt.

Microsynteny maps were constructed by aligning STORM-containing regions across species using MAFFT and comparing gene order relative to the *Scheffersomyces* species phylogeny.

Macrosynteny at the chromosomal level was assessed using ntSynt v1.03 and ntSynt-viz v1.0. We generated whole-genome alignments with ntSynt using distance thresholds -d 2 and -d 20, and normalized chromosome orientations relative to *S. shehatae* NBRC 1983 using the “normalize strand” option. Syntenic blocks containing STORM were examined for large-scale rearrangements and translocations.

### Time-calibrated species phylogeny

All focal species are represented in the calibrated time tree of Opulente, et al. (2024). To assess divergence times and affiliations of additional strains (*S. segobiensis* DSM 27193 and *S. shehatae* JCM 18690=ATY839), we estimated a time calibrated phylogeny for *Scheffersomyces*. We compiled orthologs for a panel of ten informative nuclear genes in yeast phylogenetics (Aguileta, et al. 2008; Taylor, et al. 2013), prioritizing genes present and single-copy across taxa (Supplementary Table S3). Sequences were aligned with MAFFT and concatenated. Maximum-likelihood trees were inferred in IQ-TREE 3 using a partitioned GTR+Γ model, with partitions by gene or codon position where appropriate. Divergence times were estimated in MEGA 12 using branch lengths from the ML tree and a RelTime relaxed-clock method, with internal calibrations derived from Timetree5 (Kumar, et al. 2022) using calibrations derived from TimeTree5 (Kumar et al. 2022), including the estimated *S. stipitis*-*S. lignosus* divergence of 54 MYA (range 43.2-65.1 MYA; Shen et al. 2018, 2020).

### Viral and paleoviral phylogeny

For viral and paleoviral phylogenies, we compiled amino-acid sequences of capsid domains from exogenous totiviruses, singleton totivirus-like paleoviruses, and STORM copies (TLC1-TLC4) from all sampled species. Sequences were aligned using MAFFT v7 (E-INS-i, with G-INS-i for larger datasets). To assess the effect of alignment on the results, we used ClipKIT 2.1.3 (kpic-smart-gap mode). Sequence metadata for the 84-taxon capsid amino-acid alignment underlying the main CP tree (host species, strain or isolate, primary accession, and re-parsed genomic coordinates) are provided in Supplementary Data S2. The underlying alignment (84 sequences × 956 amino-acid columns) is provided as Supplementary Data S3.

Maximum-likelihood trees were inferred in IQ-TREE 3 using ModelFinder (Crotty, et al. 2020; Wong, et al. 2025) to select the best-fitting substitution model. Branch support was assessed with approximate likelihood ratio tests (SH-aLRT) and 1,000 ultrafast bootstrap replicates. For nucleotide phylogenies of the STORM array, MixtureFinder in IQ-TREE 3 was used to identify a best-fitting heterotachy-aware mixture model. The *Scheffersomyces* TLC nucleotide alignment with two outgroups (34 sequences × 1,974 columns) underlying Figure 4 is provided as Supplementary Data S4. RdRp domains were aligned separately by the same procedure; the alignment (43 sequences × 737 columns) is provided as Supplementary Data S5 and underlies the RdRp root-to-tip comparison in Figure 6C.

Trees were rooted using a related plant virus (black raspberry virus F, EU082131.1) as an outgroup, and visualized in iTOL (Letunic and Bork 2024). Root-to-tip patristic distances were computed using Python (with Biopython).

### Tests of horizontal transfer of paleoviral arrays among yeast genomes

Incongruences between the paleoviral array and the species tree were initially assessed for support with approximate likelihood ratios. Coding sequences from *S. stipitis* CBS 6054 (GCF_000209165.1) served as tBLASTN queries against local databases of *S. stipitis* CBS 6054, *S. stipitis* NRRL Y-7124, *S. titani* CBS 13926, and *S. segobiensis* DSM 27193. After excluding capsid-like paleoviruses, orthologs 1800-2200 bp in length and present in all four taxa were retained, yielding 523 loci. Each was aligned with MAFFT and a maximum-likelihood tree estimated in IQ-TREE 3 with ModelFinder (-MFP) and 1000 ultrafast bootstraps. Patristic distances and the Mann-Whitney summary are provided in Supplementary Data S8 (stipitis_clade_strain_vs_strain); full alignments and IQ-TREE outputs are deposited on Zenodo (see Data Availability). Because STORM occupies non-syntenic positions in several species pairs, coalescent-based ILS tests are not appropriate; we instead compared patristic distances for the STORM array against the genome-wide d₁₂ distribution from single-copy reference loci using Mann-Whitney tests (patristic distances are not normally distributed). Trees for sequences flanking the putative introgressed array region (TLC1-TLC3) were estimated in IQ-TREE 3 from MAFFT alignments.

For *lignosus*/*stipitis* clade testing of HGT, a related fungus, *Lodderomyces beijingensis* (<u>GCA_963989305.1</u>) was used as an outgroup to *S. stipitis* (GCF_000209165.1), *S. lignosus* (BCGS00000000.1) and *S. xylosifermentans* (JAXBLS000000000.1). As with the *stipitis* clade, translated blasts were used to assemble orthologs. Alignments were excluded if the outgroup sequence was <500 nt, if ingroup alignments contained >50 indel gaps, or if significant ingroup base-composition differences were detected. This resulted in 272 genes. Patristic distances, the Mann-Whitney / Cliff’s-δ summary statistics, and the violin-plot summary for this comparison are provided in Supplementary Data S8 (lignosus_clade_xylo_vs_lignosus); the full alignments and IQ-TREE outputs are deposited on Zenodo (see Data Availability).

To assess the putative transfer of transposase-like elements, we initially constructed a phylogeny of IS630-like sequences in *Scheffersomyces*. The full-length protein of *S. shehatae* (BDMP01000001.1: 2358241 to 2359608) was used as a query for tblastn of *Scheffersomyces* wgs projects (save *S. spartinae* which is only distantly related to the wood-digesting species).

Fragmented pseudogenes were further reconstructed using tfasty with the Müller-Vingron VT120 matrix. A Fasta file was parsed from the hits and then aligned in MAFFT. Alignments were filtered in ClipKIT using smart-gap mode (34% of sites were filtered). Matches less than 150 AA residues were filtered. A phylogeny was estimated in IQtree3 with aLRT; model fit was estimated from the alignment. The MAFFT amino-acid alignment of 213 YEET transposases plus the *Circinella minor* IS630 outgroup (214 sequences × 950 columns) is provided as Supplementary Data S6. Terminal inverted repeats (TIRs) flanking each YEET element were identified by BLASTn searches using diagnostic 13-bp core consensus seeds (15 bp for mYEET2b). Full TIR boundaries were refined using sliding-window inverted-repeat alignments, with core sequences and parameters provided in Supplementary Data S1. For the non-autonomous mYEET2b MITE, TIR length is reported as the genome-BLAST-confirmed span of the bracketing inverted repeats (40 bp) rather than as a sliding-window score. Median aligned TIR length across confirmed elements was 191 bp (range 15-201). Element coordinates, TIR orientations, subfamily assignments, and transposase-clade membership for all 213 elements are provided in Supplementary Data S1; counts and median TIR/element sizes summarized by subfamily × clade are in Supplementary Table S2.

### Genetic distance and rate comparisons

To visualize divergence among predefined groups, pairwise patristic distances were compared using two-sided Mann-Whitney U tests with Benjamini-Hochberg correction to control the false discovery rate (FDR). These comparisons are presented as descriptive summaries of distance distributions rather than formal tests of evolutionary-rate shifts, which were instead evaluated using RELAX (see below). We quantified divergence across six groups: exogenous totiviruses, non-STORM paleoviruses, and the four STORM paralogs (TLC1-TLC4). Patristic distances were computed from maximum-likelihood trees in Python using Biopython, and leaf labels were parsed with regular expressions to assign sequences to groups. Analyses were conducted using NumPy and SciPy, and figures were generated with Matplotlib. OpenAI (GPT-5.5) was used to assist with development and debugging of custom Python scripts. The authors verified all outputs.

### Tests for selection and relaxation

We tested for shifts in selection intensity using RELAX (Wertheim et al. 2015), where K < 1 indicates relaxation and K > 1 indicates intensified selection. Significance was assessed using likelihood-ratio tests (α = 0.05). We analyzed selection intensity across STORM capsid-like genes using RELAX v4.5 in HyPhy with the Alt-Yeast-Nuclear genetic code (code 12) and a codon GTR model with three ω classes and no synonymous-rate variation. An in-frame codon alignment of 34 sequences (32 STORM genes plus two exogenous viral capsids) and the corresponding fixed phylogeny were supplied to RELAX. We ran three branch-set contrasts: (i) all STORM branches (test) versus viral capsid branches (reference); (ii) branches descending from the TLC4 paralog MRCA (test) versus all other paleoviral branches (reference), with viral branches as nuisance; and (iii) TLC1-lineage branches (test) versus TLC2/TLC3-lineage branches (reference), again with viral branches as nuisance. For each contrast we compared the K-free alternative to the K=1 null via likelihood-ratio test. Analogous RELAX analyses were applied to SKI3 orthologs to test for altered selection on this antiviral helicase in lineages with divergent STORM arrays.

To identify codons with different nonsynonymous substitution rates among capsid paralogs, we applied Contrast-FEL (Kosakovsky Pond, et al. 2021) as implemented in HyPhy (analysis version 0.61). We used the same in-frame codon alignment as for RELAX (34 taxa, 658 codons; Alt-Yeast-Nuclear genetic code) and the corresponding maximum-likelihood phylogeny. Branches leading to each tandem capsid paralog were pre-labeled in the tree according to array membership (TLC1, TLC2/3, TLC4), with viral outgroups and remaining branches treated as background. Contrast-FEL jointly estimates site-specific synonymous (α) and nonsynonymous (β) substitution rates for predefined branch sets and tests whether β differs among groups at individual codons using likelihood-ratio tests with permutation-based P-values and alignment-wide FDR correction. We conducted three pairwise contrasts (TLC1 vs TLC4, TLC2/3 vs TLC4, and TLC1 vs TLC2/3), enabling synonymous-rate variation (SRV = Yes) and permutation-based inference, and applied a nominal *P*-value threshold of 0.05 and an FDR q-value threshold of 0.2 to identify codons with significantly different dN/dS between branch sets.

### Structural prediction and comparison

Structure predictions for all capsid sequences were generated using AlphaFold3 (Abramson et al. 2024) with default parameters. All 80 ModelCIF predictions are provided in Supplementary Data S7, including STORM paralogs, TLC4 pseudogenes, singleton totivirus-like paleoviruses, Serinales-associated viral capsids, and additional totivirus references. The pLDDT scores and AlphaFold Server job IDs are tabulated in the bundled structure_inventory.tsv file (overall median pLDDT = 82.8; range 73.8-87.8). To assess whether relaxed selection affects protein structure, we computed pairwise structural similarity between paleoviral capsids and exogenous totiviral references using Foldseek (van Kempen, et al. 2024). Exogenous totivirus structures served as queries, with paleoviral structures as targets. We used qtmscore (TM-score normalized by query length) as our primary structural similarity metric because it reflects the proportion of the reference viral fold preserved in each paleoviral structure, allowing comparisons across groups with differing domain architectures.

Database creation and searching followed standard Foldseek protocols. Each paleoviral structure was compared against all exogenous viral references using a permissive E-value threshold (E ≤ 10,000) to ensure comprehensive coverage and assigned the maximum qtmscore obtained against any reference structure. Structures with no significant alignment were assigned a qtmscore of 0.0 to preserve the signal of complete structural divergence.

To focus structural comparisons on full-length capsids rather than degraded fragments, singleton paleoviral structures were post-filtered to retain only those with query coverage ≥80% when aligned to viral references. This filtering reduced the singleton dataset from 32 to 17 structures, excluding partial ORFs and highly degraded sequences while retaining all tandem array members.

Statistical comparisons of qtmscore distributions used Kruskal-Wallis tests across all groups, followed by pairwise Mann-Whitney U tests for specific hypotheses of interest. All statistical analyses were performed in Python using SciPy v1.16.1.

We mapped evolutionary rate variation and selection metrics onto the predicted structures to visualize regions of relaxed constraint and enhanced divergence. Particular attention was given to motifs implicated in decapping and RNA binding based on experimentally characterized totivirus structures (Okamoto, et al. 2020; Grybchuk, et al. 2022).

### RNA-seq analysis

Paired-end RNA-seq data for *Scheffersomyces* species under different growth conditions were obtained from NCBI SRA and JGI MycoCosm (accession numbers in Supplementary Table S4). Reads were quality-trimmed and quantified against a custom reference containing paleoviral ORFs (TLC1-TLC4), antiviral genes (SKI2, SKI3, DCP2), and housekeeping controls using Salmon v1.10.3. Expression was normalized to ACT1 TPM within each sample. Differential expression analyses used DESeq2 with lengthScaledTPM import via tximport, testing condition-specific contrasts (shake vs. baffle for *S. xylosifermentans*; chemostat vs. batch for *S. stipitis* CBS 6054) with Benjamini-Hochberg FDR correction.

## Acknowledgments

We thank Victor Albert for helpful discussions and feedback during the development of this work. We gratefully acknowledge the Center for Computational Research at the University at Buffalo for access to high-performance computing resources, and the genome centers, sequencing projects, and data submitters whose publicly available assemblies and RNA-seq datasets made this study possible.

## Data Availability

All genome assemblies and RNA-seq datasets analyzed in this study are publicly available from NCBI GenBank and the Sequence Read Archive under the accession numbers listed in the Materials and Methods. Supplementary Data S1-S6 and S8 are available as supplementary materials to this article. Custom Python and shell scripts used for patristic-distance comparisons, statistical analyses, structural-similarity compilation, and figure generation; the AlphaFold3 capsid structure predictions and Foldseek TM-score tables comprising Supplementary Data S7; and the MAFFT alignments and IQ-TREE outputs underlying the horizontal-transfer tests, are archived at Zenodo (doi:10.5281/zenodo.20261719). A portion of these data were produced by the US Department of Energy Joint Genome Institute (https://ror.org/04xm1d337; operated under Contract No. DE-AC02-05CH11231) in collaboration with the user community.

## Author Contributions

D.J.T., Conceptualization, Methodology, Formal analysis, Writing. D.A.T., Methodology, Formal analysis, Investigation, Data curation, Writing.

