## Supplementary materials for "A domesticated totivirus-like tandem array undergoes interspecific transfer and asymmetric evolution"

**This PDF file includes:** Supplementary Figures S1-S4 and Supplementary Tables S1-S6. Eight supplementary data files are provided alongside this PDF: Supplementary Data S1 (full YEET element data; Excel), S2 (Metadata for the 84-taxon capsid amino-acid alignment underlying the main CP tree; Excel), S3 (capsid amino-acid alignment; FASTA), S4 (capsid nucleotide alignment of *Scheffersomyces* TLC paralogs with two outgroups, 34 taxa × 1,974 columns; FASTA), S5 (RNA-dependent RNA polymerase amino-acid alignment, 43 taxa × 737 columns; FASTA), S6 (YEET transposase amino-acid alignment with *Circinella minor* IS630 outgroup, 214 sequences × 950 columns; FASTA), S7 (AlphaFold3 predicted capsid structures plus Foldseek TM-score tables, 80 ModelCIF files + 3 TM-score TSVs organized by capsid class; zip; archived at Zenodo, doi:10.5281/zenodo.20261719), and S8 (per-locus phylogenetic concordance summary underlying Figure 5; zip). The full reference-locus alignments and IQ-TREE outputs for both clades (~180 MB) are deposited on Zenodo and referenced in the Data Availability section.

### Supplementary Figures

**Supplementary Figure S1.** Phylogeny and structural diversification of YEET transposases in *Scheffersomyces*.

Maximum-likelihood phylogeny of YEET-family transposases identified across *Scheffersomyces* genomes, rooted with an IS630-family transposase from *Circinella minor*. Two deeply divergent transposase clades (YEET-α and YEET-β) are indicated by branch labels. Tip-associated colour strips denote terminal inverted repeat (TIR) subtype (YEET1a, YEET1b, or other), inferred from flanking sequence comparisons. Filled circles mark elements with strong TIR support, whereas filled nodes indicate branches with strong approximate likelihood-ratio test (SH-aLRT) support. Dashed connectors identify putative recent horizontal transfers. Scale bar indicates substitutions per site. Transposase coordinates, TIR sequences, and subfamily/clade assignments are presented in Supplementary Data S1.

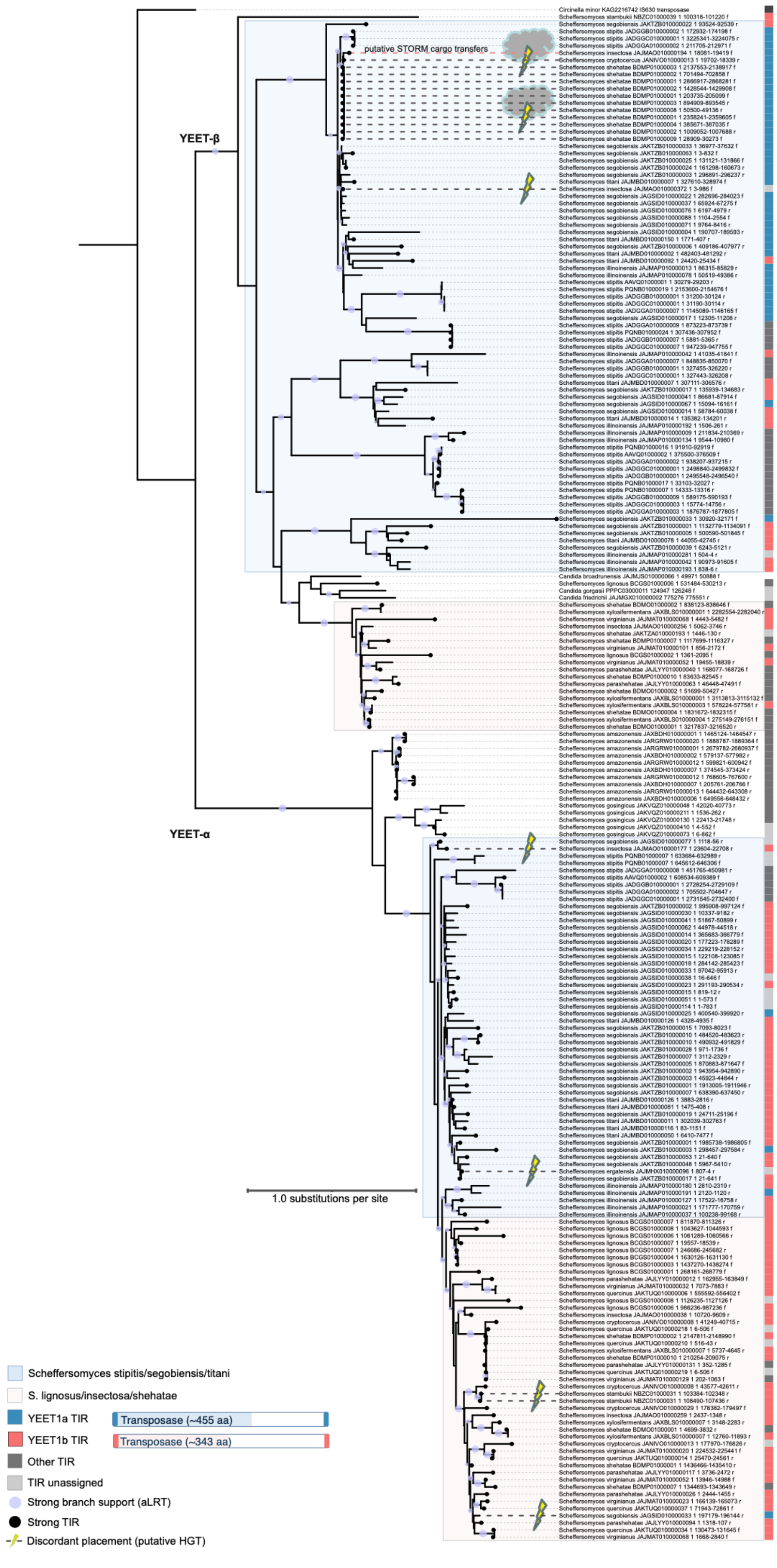
YEET-β
YEET-α
putative STORM cargo transfers
1.0 substitutions per site
Scheffersomyces stipitis/segobiensis/titani
S. lignosus/insectosa/shehatae
YEET1a TIR
Transposase (~455 aa)
YEET1b TIR
Transposase (~343 aa)
Other TIR
TIR unassigned
Strong branch support (aLRT)
Strong TIR
Discordant placement (putative HGT)

**Supplementary Figure S2.** Root-to-tip capsid (CP) distances after ClipKIT alignment trimming.

Box-and-strip plot of root-to-tip patristic distances (amino-acid substitutions per site) for the same six groups boxplotted in Figure 6A, after the MAFFT capsid amino-acid alignment was filtered with ClipKIT under the kpic-smart-gap mode (25.4% of sites removed; constant and parsimony-informative sites retained).

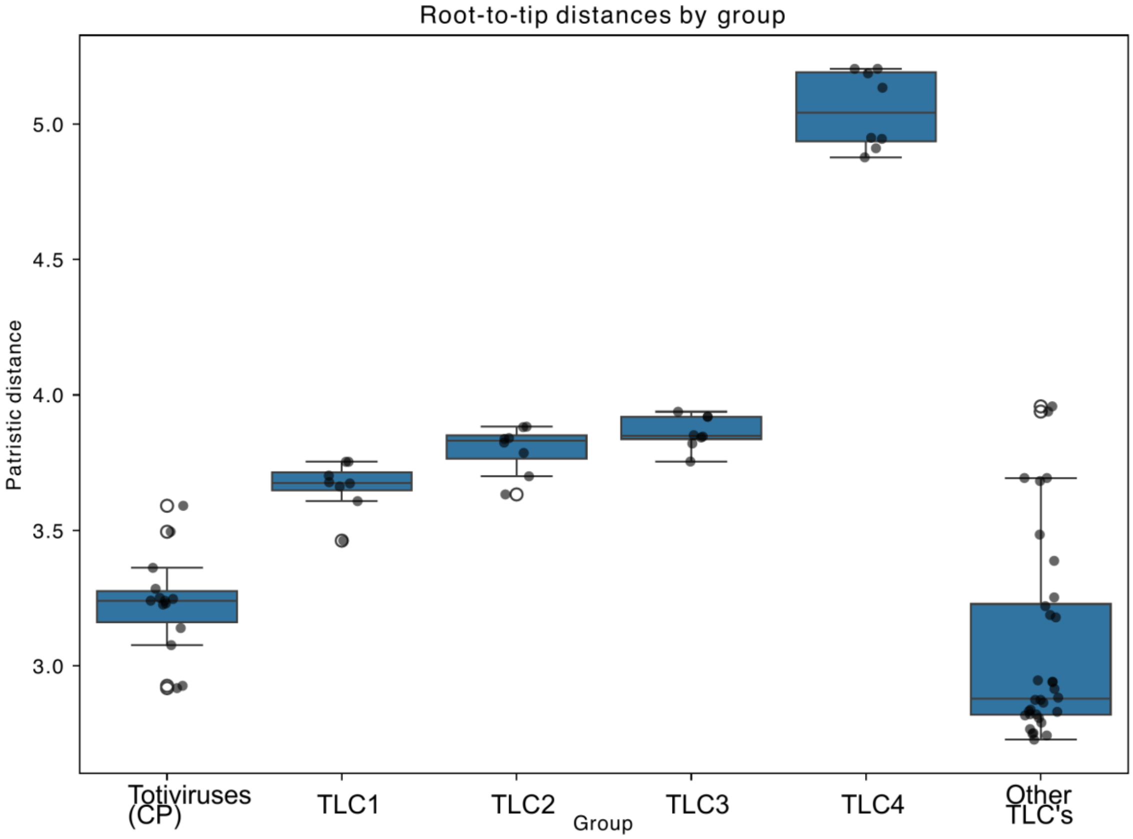

**Supplementary Figure S3.** Condition-dependent responses of STORM genes and host pathways in *Scheffersomyces*.

**(A)** Targeted differential expression in *S. xylosifermentans* comparing shake (conventional agitation) vs baffle (higher oxygen transfer and turbulence) cultures. The x-axis shows $\log_2$ fold change (shake/baffle; positive values = higher in shake), and the y-axis shows $-\log_{10}$ FDR-adjusted *P*-value (horizontal dashed line, FDR = 0.05).

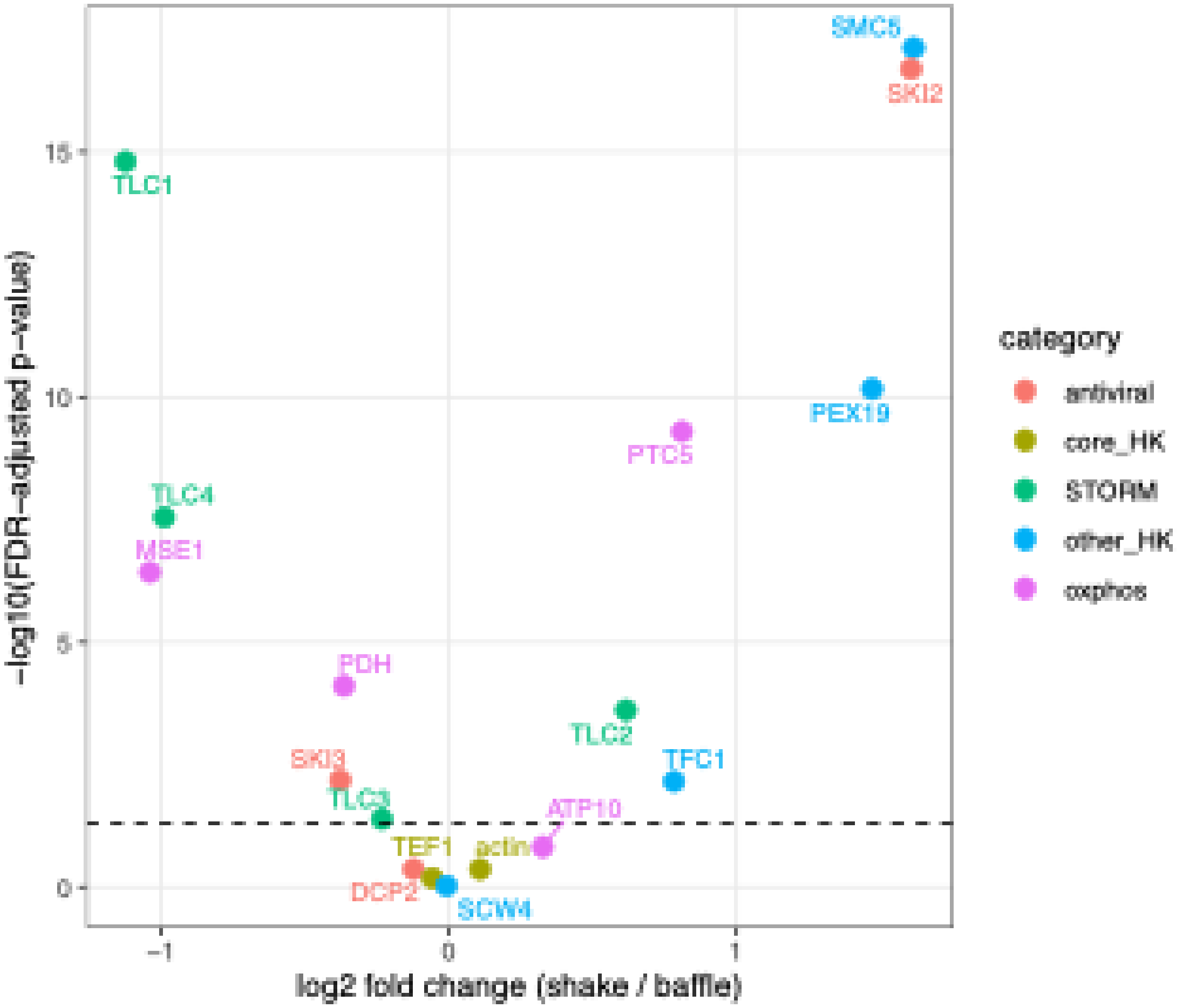

**(B)** Targeted differential expression in *S. stipitis* CBS 6054 comparing chemostat vs batch cultures ($\log_2$ fold change = chemostat/batch).

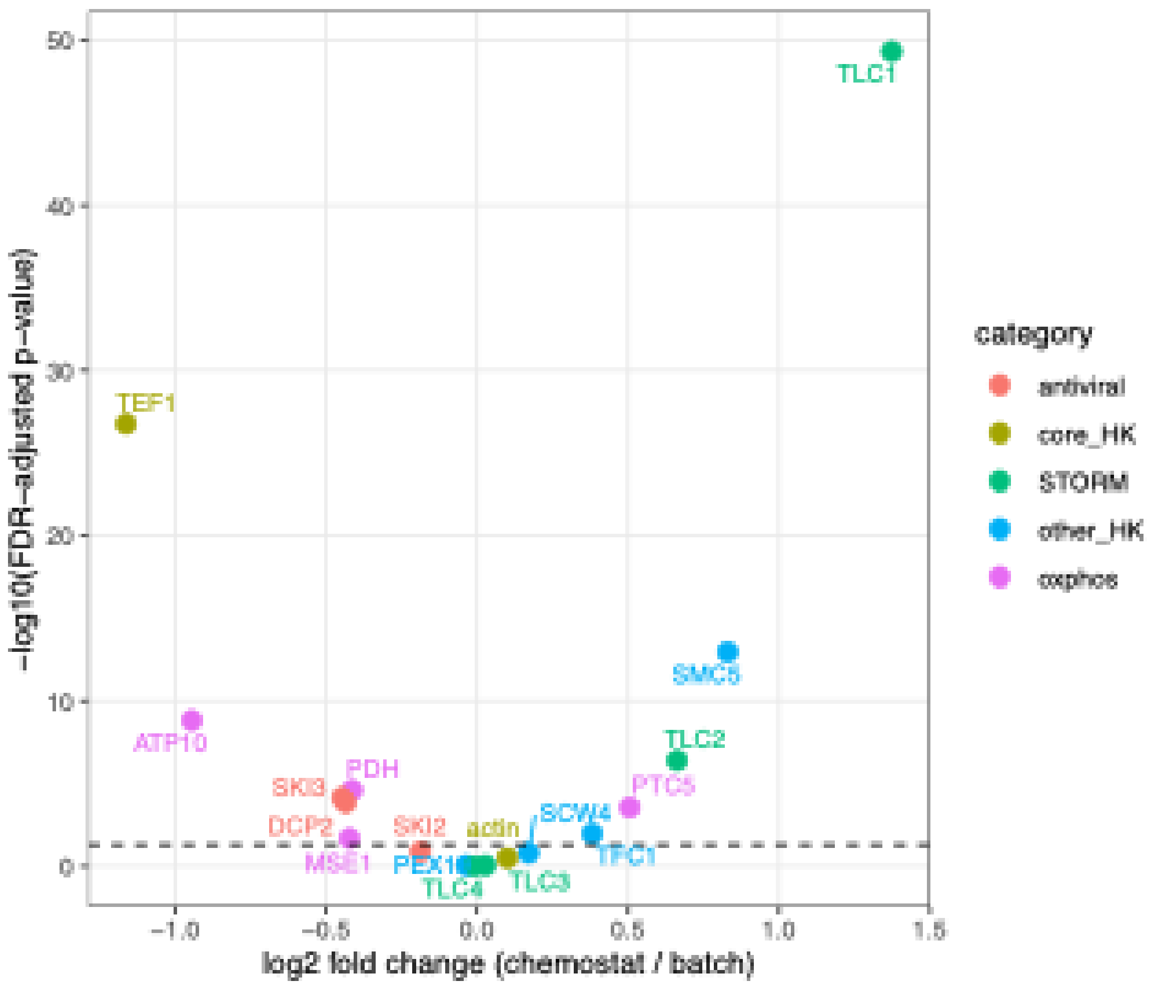

**Supplementary Figure S4.** BLAST evidence for sequence-level conservation and lineage-specific distribution of the STORM region.

BLAST hits against the *S. shehatae* ATY839 STORM-containing contig (BDMO01000001.1) ranked by maximum bit-score. Left: tabular summary (Description, Max Score, Total Score, Query Cover, E-value, Percent identity, Accession length, Accession). Right: graphical alignment of the same hits across coordinates ~1.78-1.81 Mb of the query contig. Red-shaded entries are the *shehatae*-clade *Scheffersomyces* species; blue-shaded entries are the *stipitis*-clade species, which carry STORM at a different chromosomal locus and therefore yield only short flanking-gene hits at this position.

| Description | Max Score | Total Score | Query Cover | E value | Per. ident | Acc. Len | Accession |
|---|---|---|---|---|---|---|---|
| Scheffersomyces shehatae DNA, contig: ATY839_sc01a, strain: ATY839 | 71916 | 72599 | 100% | 0.0 | 100.00 | 3488706 | BDMO01000001.1 |
| Scheffersomyces xylosifermentans strain CBS 12540 RJT20scaffold_4 | 45998 | 71114 | 100% | 0.0 | 99.25 | 1826639 | JAXBLS010000004.1 |
| Scheffersomyces lignosus DNA, scaffold: scaffold_4, strain: JCM 9837 | 5550 | 16873 | 42% | 0.0 | 83.58 | 1730773 | BCGS01000005.1 |
| Scheffersomyces shehatae DNA, contig: NBRC1983_sc05a, strain: NBRC 1983 | 5387 | 20197 | 49% | 0.0 | 83.38 | 1741095 | BDMP01000005.1 |
| Scheffersomyces quercinus strain NRRL Y-48825 NODE_6_length_736209_cov_22.039_ID_4043 | 5378 | 20144 | 47% | 0.0 | 83.39 | 736209 | JAKTUQ010000006.1 |
| Scheffersomyces cryptocercus strain NRRL Y-48824 NODE_29_length_209786_cov_64.2939_ID_6609 | 5377 | 19804 | 49% | 0.0 | 83.34 | 209721 | JANIVO010000029.1 |
| Scheffersomyces virginianus strain NRRL Y-48822 NODE_32_length_168694_cov_29.7648_ID_6328 | 5371 | 19633 | 49% | 0.0 | 83.33 | 168694 | JAJMAT010000032.1 |
| Scheffersomyces parashehatae strain CBS 12535 NODE_12_length_327702_cov_20.4121_ID_7521 | 5369 | 20141 | 49% | 0.0 | 83.31 | 327702 | JAJLYY010000012.1 |
| Scheffersomyces insectosa strain NRRL Y-12854 NODE_35_length_122011_cov_17.2874_ID_12973 | 5314 | 20358 | 51% | 0.0 | 83.04 | 122011 | JAJMAO010000035.1 |
| Scheffersomyces shehatae strain NRRL Y-12858 NODE_11_length_229527_cov_18.642_ID_7340 | 5310 | 11281 | 29% | 0.0 | 83.07 | 229527 | JAKTZA010000011.1 |
| Scheffersomyces stipitis CBS 6054 tig00000005_pilon | 1565 | 3806 | 14% | 0.0 | 81.42 | 1591760 | PQNB01000002.1 |
| Scheffersomyces titani strain CBS 13926 NODE_1_length_606302_cov_33.5475_ID_7728 | 1565 | 4067 | 21% | 0.0 | 81.42 | 606302 | JAJMBD010000001.1 |
| Scheffersomyces stipitis strain NRRL Y-7124 chromosome 3 | 1561 | 3799 | 14% | 0.0 | 81.36 | 1880751 | JADGGA010000004.1 |
| Scheffersomyces illinoinensis strain NRRL Y-48827 NODE_7_length_476829_cov_15.7456_ID_7003 | 1549 | 3776 | 15% | 0.0 | 81.05 | 476829 | JAJMAP010000007.1 |

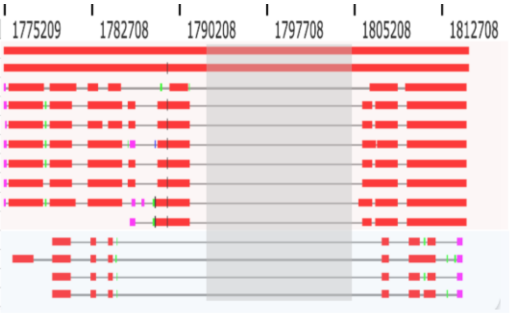

### Supplementary Tables

**Supplementary Table S1.** RELAX tests of selection intensity (Wertheim et al. 2015) on *Scheffersomyces* STORM capsid paralogs.

The RELAX test estimates a relaxation/intensification parameter *K* relative to a reference branch set, where $K < 1$ indicates relaxed purifying selection and $K > 1$ indicates intensification. LR, likelihood-ratio test statistic; *P*, associated *P*-value (df = 1). Analyses were conducted in HyPhy v2.5 using a codon alignment of 34 sequences (32 yeast paleoviruses + 2 exogenous viral capsids, 658 codons) under the Alt-Yeast-Nuclear genetic code (code 12) with a codon GTR model and three ω classes. "Other paleoviral branches" = *TLC1* + *TLC2* + *TLC3;* STORM, Scheffersomyces Totivirus-like Responsive Module; TLC, Totivirus-Like Capsid.

| Comparison | Test branch set | Reference branch set | K | LR | *P* | Interpretation |
|---|---|---|---|---|---|---|
| **STORM TLCs vs. exogenous viruses** | All STORM capsid branches | Exogenous totivirus capsid branches | 0.78 | 3.16 | 0.075 | Non-significant trend toward relaxation |
| **TLC1 vs. TLC2+TLC3** | TLC1 branches | TLC2 and TLC3 branches | 1.74 | 16.47 | **<0.001** | Significant intensification of purifying selection on TLC1 |
| **TLC4 vs. other STORM TLCs** | TLC4 branches | All other paleoviral capsid branches | 0.29 | 74.12 | **<0.001** | Strong, significant relaxation of selection on TLC4 |

**Supplementary Table S2.** YEET element counts by TIR subfamily and transposase clade.

Counts of 213 YEET elements broken down by TIR-sequence-based subfamily (rows) and transposase-phylogeny-based clade (columns). mYEET2b is a non-autonomous MITE (miniature inverted-repeat transposable element) derived from the YEET2b lineage; ? indicates tentative (TIR only) assignment of subfamily. Data are provided in Supplementary Data S1.

| TIR subfamily | YEET-α (n) | YEET-β (n) | Total (n) | Median TIR length (bp) | Median element size (bp) | n with TIR / n with size |
|---|---|---|---|---|---|---|
| YEET1a | 4 | 41 | 45 | 174 | 1809 | 17 / 34 |
| YEET1b | 72 | 23 | 95 | 188 | 1524 | 45 / 80 |
| YEET1b-Tpase | 2 | 0 | 2 | NA | NA | 0 / 0 |
| YEET1c | 5 | 8 | 13 | 174 | 2225 | 6 / 13 |
| YEET1c? | 1 | 0 | 1 | 201 | 1596 | 1 / 1 |
| YEET1d | 0 | 2 | 2 | 184 | 1577 | 2 / 2 |
| YEET1d-like? | 0 | 1 | 1 | NA | NA | 0 / 0 |
| YEET1e | 2 | 10 | 12 | 195 | 1791 | 12 / 12 |
| YEET1f | 1 | 7 | 8 | 194 | 1802 | 8 / 8 |
| YEET1f-Tpase | 0 | 4 | 4 | NA | NA | 0 / 0 |
| YEET1h? | 0 | 1 | 1 | NA | 3430 | 0 / 1 |
| YEET2a | 10 | 0 | 10 | 201 | 1369 | 10 / 10 |
| YEET2b | 1 | 0 | 1 | 201 | 1941 | 1 / 1 |
| mYEET2b † | 2 | 0 | 2 | 40 | 1719 | 1 / 2 |
| YEET-β-Candida | 0 | 1 | 1 | NA | 2216 | 0 / 1 |
| unassigned | 12 | 3 | 15 | NA | 1535 | 0 / 5 |
| **Total** | **112** | **101** | **213** | | | |

**Supplementary Table S3.** Reference nuclear genes used for species-tree inference and introgression testing in *Scheffersomyces*.

Accession numbers are from *S. stipitis* CBS 6054 (GCF_000209165.1) except KC616421.1 (*ELP3*). Gene product annotations are from UniProt/SGD. The same gene panel was used in Aguileta et al. (2008) and Taylor et al. (2013).

| Accession | Gene | Gene product / function |
|---|---|---|
| XM_001384142.1 | ***MCM7*** | Mini-chromosome maintenance complex component 7 |
| KC616421.1 | ***ELP3*** | Elongator complex protein 3 |
| XM_001387404.1 | ***KOG1*** | TORC1 subunit KOG1 (Raptor homolog) |
| XM_001384848.1 | ***GDH2*** | NAD-dependent glutamate dehydrogenase |
| XM_001384577.1 | ***ILV2*** | Acetolactate synthase, large subunit |
| XM_001382598.1 | ***UBA1*** | Ubiquitin-activating enzyme E1 1 |
| XM_001386823.1 | ***POL1*** | DNA polymerase alpha, catalytic subunit |
| XM_001387653.1 | ***RBV1*** | Protein RBV1 (Ribosome biogenesis) |
| XM_001384154.1 | ***MSH3*** | DNA mismatch repair protein MSH3 |
| XM_001387301.1 | ***CCT8*** | T-complex protein 1 subunit theta (CCT8) |

**Supplementary Table S4.** Public RNA-seq datasets used for targeted STORM expression analyses.

Paired-end RNA-seq datasets for Scheffersomyces species under different growth conditions, obtained from the NCBI Sequence Read Archive (SRA) and JGI MycoCosm. Conditions correspond to those analysed in Figure 10 and Supplementary Figure S3 (shake vs baffle for *S. xylosifermentans* CBS 12540; aerobic batch vs aerobic chemostat for *S. stipitis* CBS 6054).

| Species | Strain / isolate | Condition contrast | Public source(s) and accession(s) | Read format | Notes |
|---|---|---|---|---|---|
| *Scheffersomyces xylosifermentans* | CBS 12540 | Moderate aeration (shake flask) vs high aeration (baffled flask). Culture: 125-mL unbaffled vs 250-mL baffled flasks, 50 mL YPX (5% xylose), 30 °C, 200 rpm, 72 h. | NCBI SRA BioProject PRJNA1015965; JGI MycoCosm portal Schxyl1; JGI Project ID 1175645. Condition-specific SRX/SRR run IDs to be filled from the SRA Run Selector before submission. | Illumina NovaSeq S4, paired-end 2 × 151 bp | Barros et al. (2024); shake-vs-baffle contrast used for Figure 10 and Supplementary Figure S3 (panel A). |
| *Scheffersomyces stipitis* | CBS 6054 | Aerobic batch vs aerobic chemostat. Batch on 20 g/L glucose; chemostat on 2 g/L glucose at D = 0.1 $h^{-1}$. RNA sampled at mid-exponential phase ($OD_{600}$ = 2.5). | NCBI SRA Study SRP012047; BioSample SAMN00849586; Sample SRS308058; Experiment SRX135712; Runs SRR453575, SRR453576, SRR453577. | Illumina Genome Analyzer, poly(A)-selected paired-end 2 × 102 bp | Papini et al. (2012); batch-vs-chemostat contrast used for Figure 10 and Supplementary Figure S3 (panel B). Indexed SRA page is titled "Chemostat cultivation S. stipitis"; the source paper describes the comparison as batch vs chemostat. |

**Supplementary Table S5.** Group descriptive statistics for capsid root-to-tip patristic distances (Figure 6A).

Mean and standard deviation of root-to-tip distances (expected amino-acid substitutions per site) computed from the maximum-likelihood capsid amino-acid tree underlying Figure 6A. Group identities (1-6) correspond to the boxplot categories in Figure 6A. Pairwise tests of these distributions are reported in Supplementary Table S6.

| Group | Identity | n | Mean root-to-tip | SD |
|---|---|---|---|---|
| **1** | Exogenous viruses | 14 | 3.23 | 0.185 |
| **2** | STORM TLC1 | 8 | 3.661 | 0.094 |
| **3** | STORM TLC2 | 8 | 3.798 | 0.089 |
| **4** | STORM TLC3 | 8 | 3.862 | 0.062 |
| **5** | STORM TLC4 | 8 | 5.051 | 0.143 |
| **6** | Non-STORM paleoviruses (Other NIRVs) | 32 | 3.069 | 0.372 |

**Supplementary Table S6.** Pairwise Mann-Whitney U tests of capsid patristic distances between the groups listed in Supplementary Table S5.

Distances were extracted from the maximum-likelihood capsid amino-acid phylogeny (see Methods). Two-sided U statistics and raw *P*-values are reported alongside Benjamini-Hochberg FDR-corrected *P*-values across all 15 comparisons. Sample sizes: exogenous viruses (n = 14), TLC1-TLC4 (n = 8 each), Other NIRVs (non-STORM totivirus-like capsid paleoviruses; n = 32).

| Group A | Group B | U | P (raw) | P (FDR) | Sig. |
|---|---|---|---|---|---|
| **Exogenous viruses** | **TLC1** | 2 | 2.50e-05 | **9.38e-05** | **Yes** |
| **Exogenous viruses** | **TLC2** | 0 | 6.25e-06 | **4.69e-05** | **Yes** |
| **Exogenous viruses** | **TLC3** | 0 | 1.51e-04 | **3.05e-04** | **Yes** |
| **Exogenous viruses** | **TLC4** | 0 | 6.25e-06 | **4.69e-05** | **Yes** |
| **Exogenous viruses** | **Other NIRVs** | 326 | 0.0154 | **0.0165** | **Yes** |
| **TLC1** | **TLC2** | 9 | 0.0148 | **0.0165** | **Yes** |
| **TLC1** | **TLC3** | 0 | 9.31e-04 | **1.27e-03** | **Yes** |
| **TLC1** | **TLC4** | 0 | 1.55e-04 | **3.05e-04** | **Yes** |
| **TLC1** | **Other NIRVs** | 224 | 1.24e-03 | **1.55e-03** | **Yes** |
| **TLC2** | **TLC3** | 17 | 0.128 | 0.128 | *n.s.* |
| **TLC2** | **TLC4** | 0 | 1.55e-04 | **3.05e-04** | **Yes** |
| **TLC2** | **Other NIRVs** | 237 | 2.43e-04 | **4.06e-04** | **Yes** |
| **TLC3** | **TLC4** | 0 | 9.31e-04 | **1.27e-03** | **Yes** |
| **TLC3** | **Other NIRVs** | 240 | 1.63e-04 | **3.05e-04** | **Yes** |
| **TLC4** | **Other NIRVs** | 256 | 1.62e-05 | **8.10e-05** | **Yes** |

### Supplementary Data

**Supplementary Data S1.** YEET (Yeast Endogenous Element Transposon) element data file :Supplementary_Data_S1_YEET_elements.xlsx.

**Supplementary Data S2.** Metadata for the 84-taxon capsid (CP) amino-acid alignment underlying the main CP tree (Figures 3 and 6A): CP_MAFFT_84_taxa_supplement.xlsx.

**Supplementary Data S3.** Capsid (CP) amino-acid alignment underlying the main CP tree (Figures 3 and 6A).

**Supplementary Data S4.** Capsid (CP) nucleotide alignment of *Scheffersomyces* TLC paralogs with two outgroups (Figure 4).

**Supplementary Data S5.** RNA-dependent RNA polymerase (RdRp) amino-acid alignment (Figure 6C). The alignment underlies the RdRp root-to-tip distance comparison in Figure 6C.

**Supplementary Data S6.** YEET transposase amino-acid alignment (used for Supplementary Figure S1).

**Supplementary Data S7.** AlphaFold3 predicted capsid structures underlying Figures 7 and 8.

Zip archive (Supplementary_Data_S7_AlphaFold3_predictions.zip; ~8 MB compressed), archived at Zenodo (doi:10.5281/zenodo.20261719).

**Supplementary Data S8.** Phylogenetic concordance summary underlying Figure 5.

MAFFT alignments and IQ-TREE outputs (~180 MB across both clades) are deposited on Zenodo (see Data Availability for the DOI).
