## Supplementary material for "A domesticated totivirus-like tandem array undergoes interspecific transfer and asymmetric evolution": HGT_vs_Species_stats_summary.pdf

### HGT vs Species — Exact/Permutation MWU and Cliff's δ

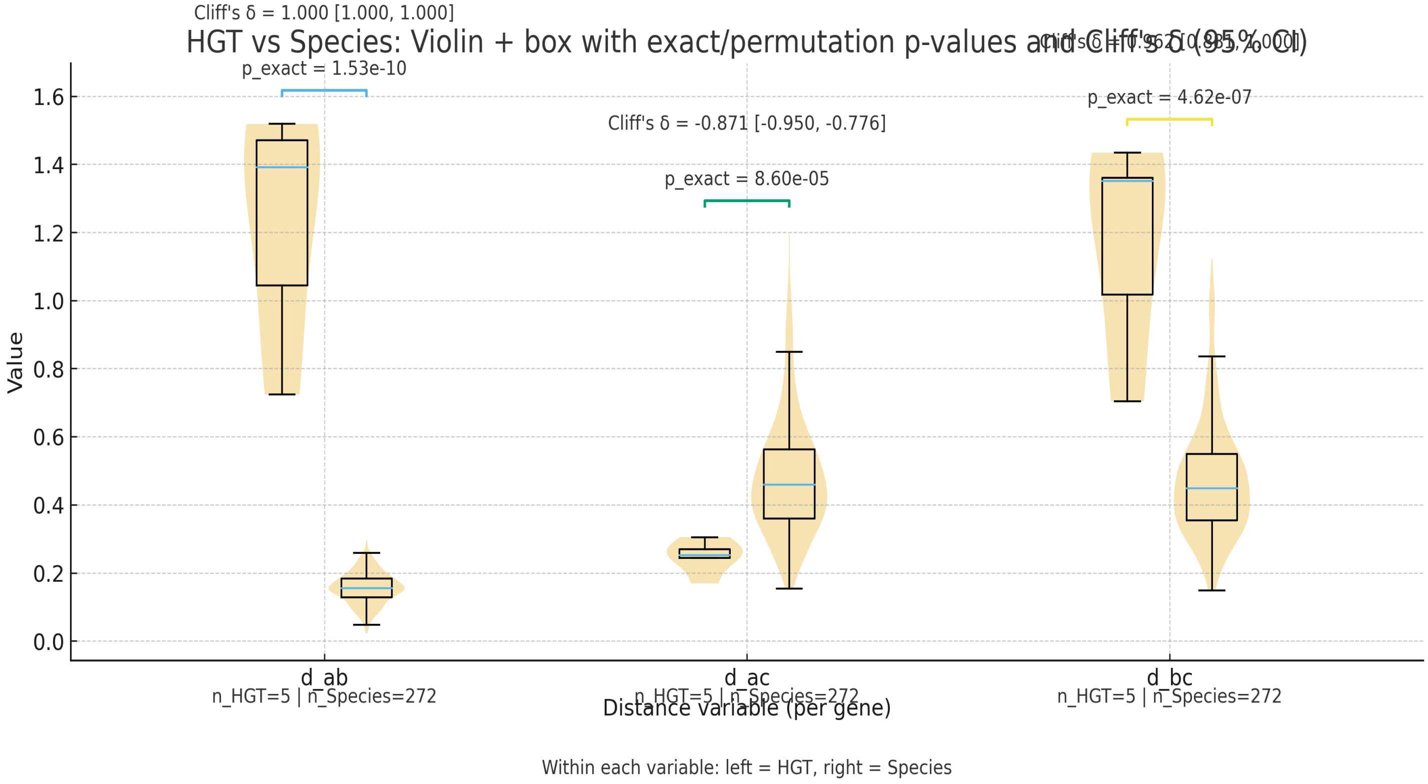


| Variable | n_HGT | n_Species | U_stat | p_exact | p_perm | Cliff's δ | δ_lo | δ_hi |
|---|---|---|---|---|---|---|---|---|
| d_ab | 5 | 272 | 1360.0 | 1.526e-10 | 5.000e-05 | 1.000 | 1.000 | 1.000 |
| d_ac | 5 | 272 | 88.0 | 8.597e-05 | 1.500e-04 | -0.871 | -0.950 | -0.776 |
| d_bc | 5 | 272 | 1334.0 | 4.622e-07 | 5.000e-05 | 0.962 | 0.881 | 1.000 |
